# DeepsmirUD: Precise prediction of regulatory effects on miRNA expression mediated by small molecular compounds using competing deep learning frameworks

**DOI:** 10.1101/2022.06.30.498304

**Authors:** Jianfeng Sun, Jinlong Ru, Zihao Chen, Fei Qi, Lorenzo Ramos-Mucci, Suyuan Chen, Adam P. Cribbs, Li Deng, Xia Wang

**Author notes:** These authors contributed equally to this work (Jianfeng Sun, Jinlong Ru).

## Abstract

Aberrant miRNA expression has pervasively been found to relate to a growing number of human diseases. Therefore, targeting miRNAs to regulate their expression levels has become an important therapy against diseases that stem from the dysfunction of oncogenic pathways regulated by the miRNAs. In recent years, small molecule compounds have demonstrated enormous potential as drugs to regulate miRNA expression (*i*.*e*., SM-miR). A clear understanding of the mechanism of action of small molecules on down- and up-regulating miRNA expression allows precise diagnosis and treatment of oncogenic pathways. However, outside of a slow and costly process of experimental determination, computational strategies to assist this in an ad hoc manner have still not been enabled. In this work, we develop, to the best of our knowledge, the first prediction tool, DeepsmirUD, to infer small molecule-mediated regulatory effects on miRNA expression. This method is powered by an ensemble of 12 cutting-edged deep learning frameworks and achieves state-of-the-art performance with AUC values of 0.840/0.969 and AUCPR values of 0.866/0.983 on two independent test datasets. With a complementarily constructed network inference approach based on similarity, we report a significantly improved accuracy of 0.813 in determining regulatory effects of nearly 650 SM-miR relations formed with either novel small molecules or novel miRNAs. By further integrating miRNA-cancer relations, we established a database of potentially pharmaceutical drugs to aid in understanding the drug mechanisms of action in diseases and to offer novel insight into drug repositioning. Taken together, our method shows great promise to assist and accelerate the therapeutic development of potential miRNA targets and small molecule drugs. Furthermore, we have employed DeepsmirUD to predict regulatory effects of a large number of high-confidence SM-miR relations curated from Psmir, which are publicly available through https://github.com/2003100127/deepsmirud and https://rujinlong.github.io/deepsmirud/ alongside the DeepsmirUD standalone package.

## Introduction

microRNAs (miRNAs) are a class of non-coding RNAs of approximately 20nt in size [1], which are vital to biological activities by primarily virtue of their post-transcriptionally regulatory mechanisms [2,3]. miRNAs are often known to downregulate the expression of genes by inhibiting translation or promoting the degradation of target mRNAs [4,5], thereby exerting an impact on the gene regulatory pathways to either remodel homeostasis [6] or give rise to malignancy [7] such as leukemia [8] and osteoarthritis [9]. It has been reported that up to thousands of mRNAs can be targeted by a single miRNA and vice versa [10]. Therefore, there are a myriad of gateways for miRNAs to influence the regulated gene pathways. Growing evidence has particularly suggested that miRNAs play important roles in cancer [11] and their expression abnormalities can lead to a variety of diseases [12,13] or supress tumor progression [14,15]. For example, suppression of apoptosis in *Myc*-induced lymphomas entails amplifying the miR-17/92 cluster, whereas the genetic ablation of the endogenous miR-17/92 allele can enhance apoptosis and reduce tumorigenicity. Furthermore, downregulated *let-7* miRNA expression levels are widely detected in patients with Burkitt lymphoma [16], whereas overexpression of *let-7* miRNAs is found to repress tumorigenesis by upregulating the expression of RAS-encoded transmembrane proteins to inhibit cellular growth and differentiation [11]. In this regard, therapeutics can be achieved by targeting oncogenic miRNAs with potential drug molecules to alter their expression [17–20]. Existing experiments have revealed that many small molecules (SMs) have held great promise as pharmaceuticals to target the miRNAs to modulate their expression thereafter [21–24].

Experimentally verified measures to determine whether binding relationships exist between small molecules and miRNAs (SM-miR) are normally time-consuming and costly [25,26] in that it is intractable to experimentally study all possible binary combinations giving a pool of SM and miRNA candidates. According to the SM2miR database built based on more than 2000 publications [27], only 1.14% of all possible SM-miR pairs interwoven with 1492 unique miRNAs and 212 unique small molecules (after pre-processing) are catalogued to be experimentally verified. It is therefore in addition to the experimental profiling that much attention has been paid for designing computational techniques to speed up the inference of binding. Over the past decade, approaches for predicting the binding of small molecules to miRNAs have been extensively studied and the predictive power has been gradually sharpened [28–30]. This has led to methods developed especially based upon similarity (e.g., EKRRSMMA [31] and BNNRSMMA [32]) or machine-learning inference techniques (e.g., ELMDA [33] and PSRR [34]). Despite tremendous effort made in determining whether small molecules bind to miRNAs, questions like in which regulation types (*i*.*e*., downregulation or upregulation) the miRNA expression is mediated by the binding small molecules and how likely and strongly they downregulate or upregulate the expression have remained computationally unexplored in an ad hoc manner. A precise understanding of the types of SM-mediated regulation of miRNA expression may provide direct or indirect evidence for cancer pathogenesis and therapeutics in the sense that the oncogenic signalling pathways regulated by the miRNAs are affected. Therefore, the efficient identification and the follow-up analysis of disease-associated miRNAs targeted by small molecule drugs with a measurable influence on the miRNA expression are becoming increasingly important [22,30,35] and have emerged as a new therapeutic treatment in miRNA pharmacogenomics [36].

Advances in deep learning have spawned ample opportunity to promote biological applications and discoveries [37] such as protein structural [38] and functional [39] prediction. The centrality of maintaining and stepping the advances forward has so far been the adoption of two chief neural layers, convolutional and recurrent neural layers, which are primarily used as basic structures in designing computer-vision and speech-recognition methods, respectively. Massive variant deep learning architectures can be devised through lessening or heightening the complexity in and/or between layers [40]. These have led to remarkable progress in powering and improving biomedical findings [41–43]. In order for maximization of method performance specific to the regulatory effect prediction, we attempted to seek prime solutions from the computer-vision and speech-recognition fields by utilizing both well-established models including AlexNet [44], MobileNetV2 [45], Transformer-based ConvMixer [46], ResNet18 [47], ResNet50 [47], attention-based SCAResNet18 [48], and our self-assembly architectures, including convolutional neural networks (CNNs) [49], recurrent neural networks (RNNs) [50], bidirectional recurrent neural networks (BiRNNs) [51], depth-wise and separable neural networks [52], the fusion of long short-term memory (LSTM) neural networks and CNNs [53,54], and sequence-to-sequence (Seq2Seq) neural networks [55,56]. These models, the vast majority of which can be trained fast due to the residual connection in design [47] or required parameter numbers, allow relatively full-scale examination and comparison of performance from shallow to ultradeep layers visually and semantically.

Here, we present the first construction of a deep learning niche comprising 12 competing frameworks to predict SM-mediated regulatory effects upon miRNA expression. A comprehensive analysis was made to opt for deep learning models optimized sufficiently and properly based on well-curated SM-miR relations and biophysical and biochemical features. Through avoiding overtraining rigorously, LSTMCNN is reported to outperform the rest of models on experimentally resolved relations, and ResNet-based models, including ResNet18, ResNet50, and SCAResNet18, are preferable to attain the most stable prediction performance over long training epochs. In the context of achieving competing results (at least AUC 0.80-0.92) by individual models, the final ensemble model, DeepsmirUD, can tap into their variance-reduced predictions to obtain an even further boosted performance gain up to ∼2% in AUC and ∼1-2% in AUCPR. A network inference approach is also added to this new tool, which enables significant improvement for SM-miR relations of novel miRNA targets or SM drugs. By trawling through miRNA-disease data from miRCancer [57], we finally established a database of drug–cancer associations to provide potential therapeutics based on the SM-miR upregulation and downregulation profiles predicted by DeepsmirUD.

## Results

### Model determination upon full-scale stabilities in predicting regulatory effects

Determination of models better trained over epochs is one of the major precursors of maximizing predictive performance when the models are optimized on a dataset of finite size. Often, limited by the size of collected samples, models do not have to necessarily perform ideally all along on all kinds of test data. However, if quality control of the models during/after training was poorly conducted, their generalization abilities on different kinds of test data could become even worse, thereby likely losing magnitude of the models to be used in real-world applications. It was shown in a previous study that the performance of a model could slip grossly when it was examined on different test datasets [58–60]. To manage to secure models with a better generalization ability, we enlarged the size of data required for training, mixed the SM-miR relations from different sources, employed a wide array of deep learning algorithms, and fully monitored performance variations via various assessment metrics with a large epoch number (**Figure 3**). Apart from MobileNetV2, the rest of deep learning approaches mostly present a quite stable performance variation on the ebb and flow basis across 400 epoches. After reaching the AUC value of 0.8 in dozens of epochs, around half of all methods have levelled off as evidenced by the overall profiles and statistically significant p-values and R-square coefficients (i.e., >0.8). In combinations of AUC and AUCPR profiles, ResNet-wise methods, including ResNet18, ResNet50, and SCAResNet18, are more prone to obtain the most stable performance, while MobileNetV2 and the up-to-date ConvMixer64 method are more vulnerable to the Test dataset with oscillation between AUC values 0.6 and 0.8 (**Figures 3a and 3c**). Apparently, these two methods are not recommendations for applications in modelling regulatory effects. In addition, BiRNN shows a decrease in AUC over the epochs. Overall, the majority of the deep learning approaches carve out steady performance, which provides proof that the curated features used could be better acquired by the deep learning frameworks. The best models for each method were finally determined by the early stopping strategy. Moveover, AUCPR, ACC, bACC, precision, recall, F1-score, MCC, Jaccard values changing with epochs on the Test and TestSim datasets can be found in **Supplementary Figures S1-S16**.

**Figure 1.**
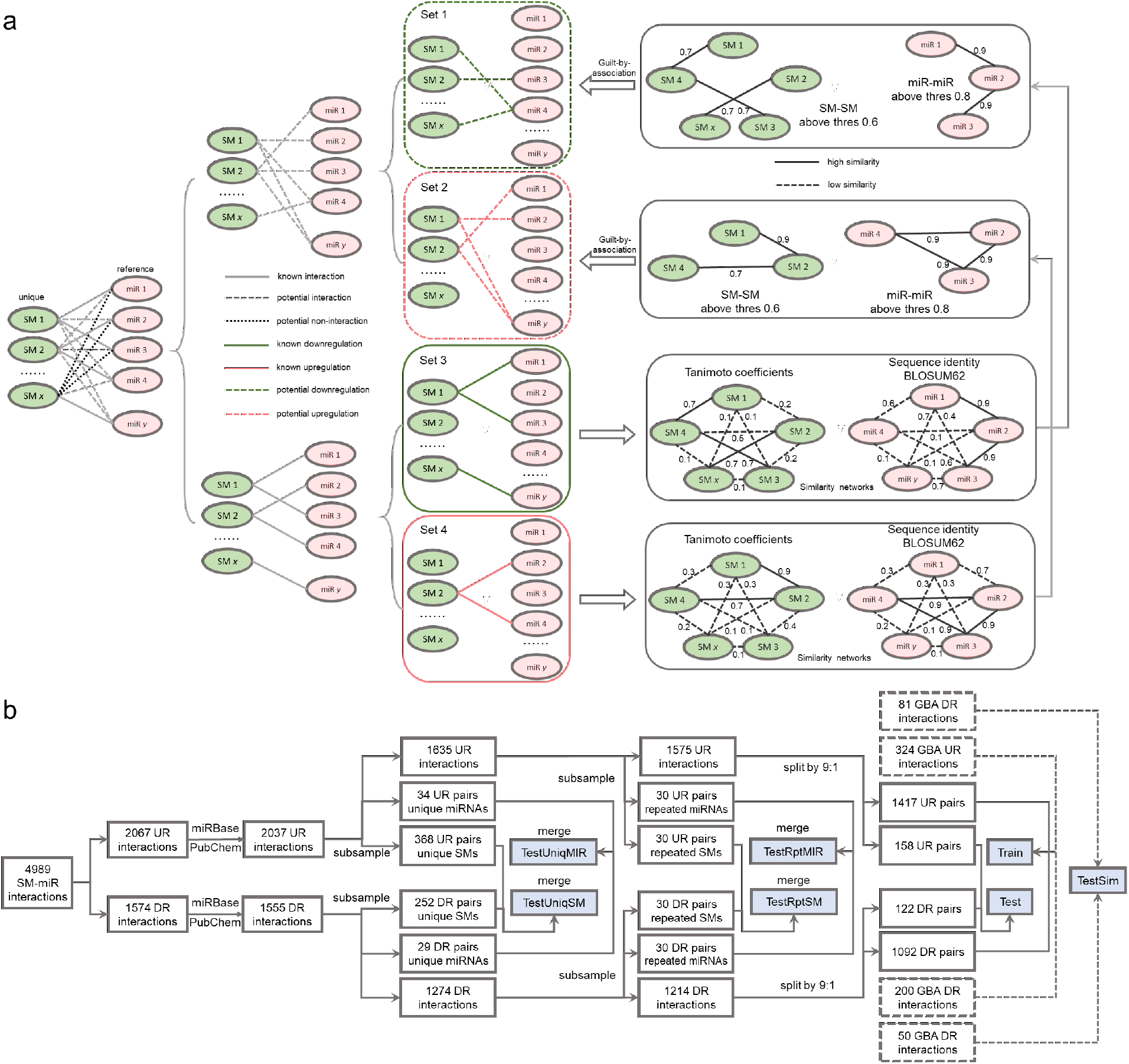
Data wrangling of upregulated and downregulated relations. a. Inference of potentially upregulated and downregulated relations using guilt-by-association. b. Flowchart of generating the Train, Test, TestSim, TestRptMIR, TestRptSM, TestUniqMIR, and TestUniqSM datasets.

**Figure 2.**
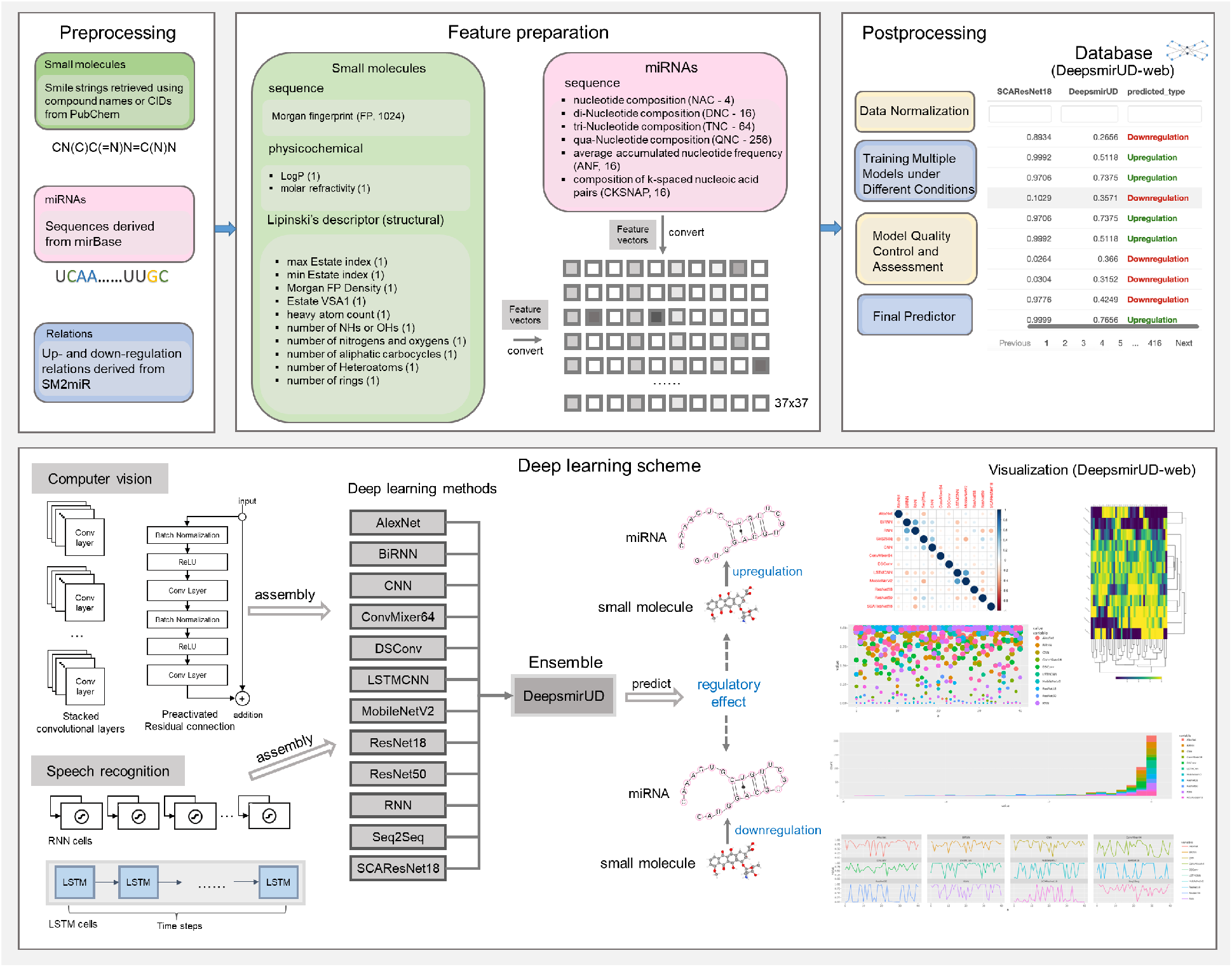
Workflow of predicting SM-mediated regulatory effects on miRNA expression by deep learning algorithms. In box *Feature preparation*, the integers stand for the length of features. The feature vector stands for a concatenation of miRNA and small-molecule features.

**Figure 3.**
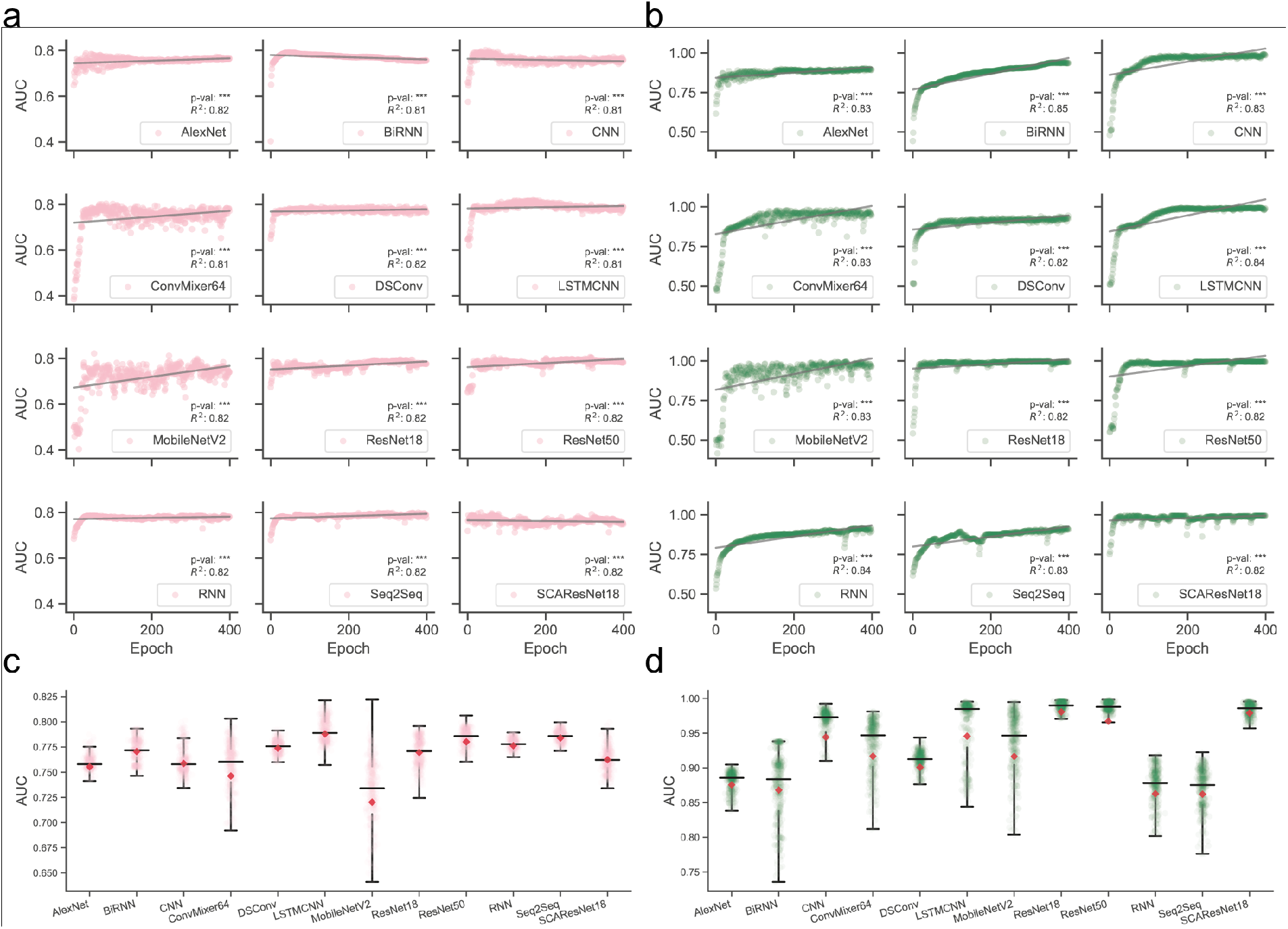
Overview of full-scale performance examination of deep learning algorithms on independent test SM-miR relations over training epochs. Landscapes of AUC variations on Test (a) and TestSim (b). Boxplots of AUC values on Test (c) and TestSim (d).

### Performance of SM-mediated regulatory effects on miRNA expression using independent SM-miR relations

The models produced by all individual methods and determined using the above procedures have exhibited high performance in predicting regulatory effects (**Figure 4, Table 1, Supplementary Figure S17a, and Supplementary Table S1**). Among them, LSTMCNN shows the best AUC and AUCPR performance. As a result of integrating all 12 models together, the ensemble model, DeepsmirUD, outperforms all of them with an AUC value 0.840 and an AUCPR value 0.866 on the Test dataset. It shows a very similar profile with LSTMCNN (**Figure 4c**). All models achieve at least above AUC 0.770 and AUCPR 0.810. The ResNet-based methods give a quite even distribution of prediction values between downregulation and upregulation classes, while RNN-based methods mostly yield prediction values floating around 0.5 (**Figure 4c and 4d**). This may suggest that RNN-based methods do not well learn the characterization of regulation types and so the values compromise in the middle. In addition, ConvMixer64, one of the most recent Vision Transformer (ViT) methods using patch learning [46], do not give accurate results by its one-sided prediction values (**Figure 4d**) but obtain the best precision of over 0.8 (**Figure 4b**). Almost all methods have an insignificant difference in ACC and F1-score.

**Table 1.** Prediction performance evaluation on Test.

| Method | AUC | AUCPR | ACC | bACC | Precision | Recall | MCC | F1score | Jaccard |
| --- | --- | --- | --- | --- | --- | --- | --- | --- | --- |
| AlexNet | 0.779 | 0.827 | 0.691 | 0.688 | 0.730 | 0.712 | 0.374 | 0.721 | 0.563 |
| BiRNN | 0.790 | 0.828 | 0.691 | 0.681 | 0.708 | 0.763 | 0.367 | 0.735 | 0.580 |
| RNN | 0.782 | 0.817 | 0.712 | 0.704 | 0.732 | 0.769 | 0.412 | 0.750 | 0.600 |
| Seq2Seq | 0.796 | 0.824 | 0.698 | 0.687 | 0.712 | 0.776 | 0.381 | 0.742 | 0.590 |
| CNN | 0.796 | 0.829 | 0.723 | 0.710 | 0.726 | 0.814 | 0.432 | 0.767 | 0.623 |
| ConvMixer64 | 0.804 | 0.819 | 0.691 | 0.711 | 0.850 | 0.545 | 0.436 | 0.664 | 0.497 |
| DSConv | 0.791 | 0.828 | 0.687 | 0.663 | 0.673 | 0.859 | 0.359 | 0.755 | 0.606 |
| LSTMCNN | 0.822 | 0.848 | 0.759 | 0.749 | 0.760 | 0.833 | 0.507 | 0.795 | 0.660 |
| MobileNetV2 | 0.781 | 0.810 | 0.723 | 0.726 | 0.780 | 0.705 | 0.448 | 0.741 | 0.588 |
| ResNet18 | 0.789 | 0.838 | 0.701 | 0.704 | 0.759 | 0.686 | 0.404 | 0.721 | 0.563 |
| ResNet50 | 0.804 | 0.820 | 0.730 | 0.726 | 0.758 | 0.763 | 0.452 | 0.760 | 0.613 |
| SCAResNet18 | 0.792 | 0.821 | 0.723 | 0.713 | 0.734 | 0.795 | 0.433 | 0.763 | 0.617 |
| DeepsmirUD | 0.840 | 0.866 | 0.763 | 0.756 | 0.778 | 0.808 | 0.516 | 0.792 | 0.656 |

**Figure 4.**
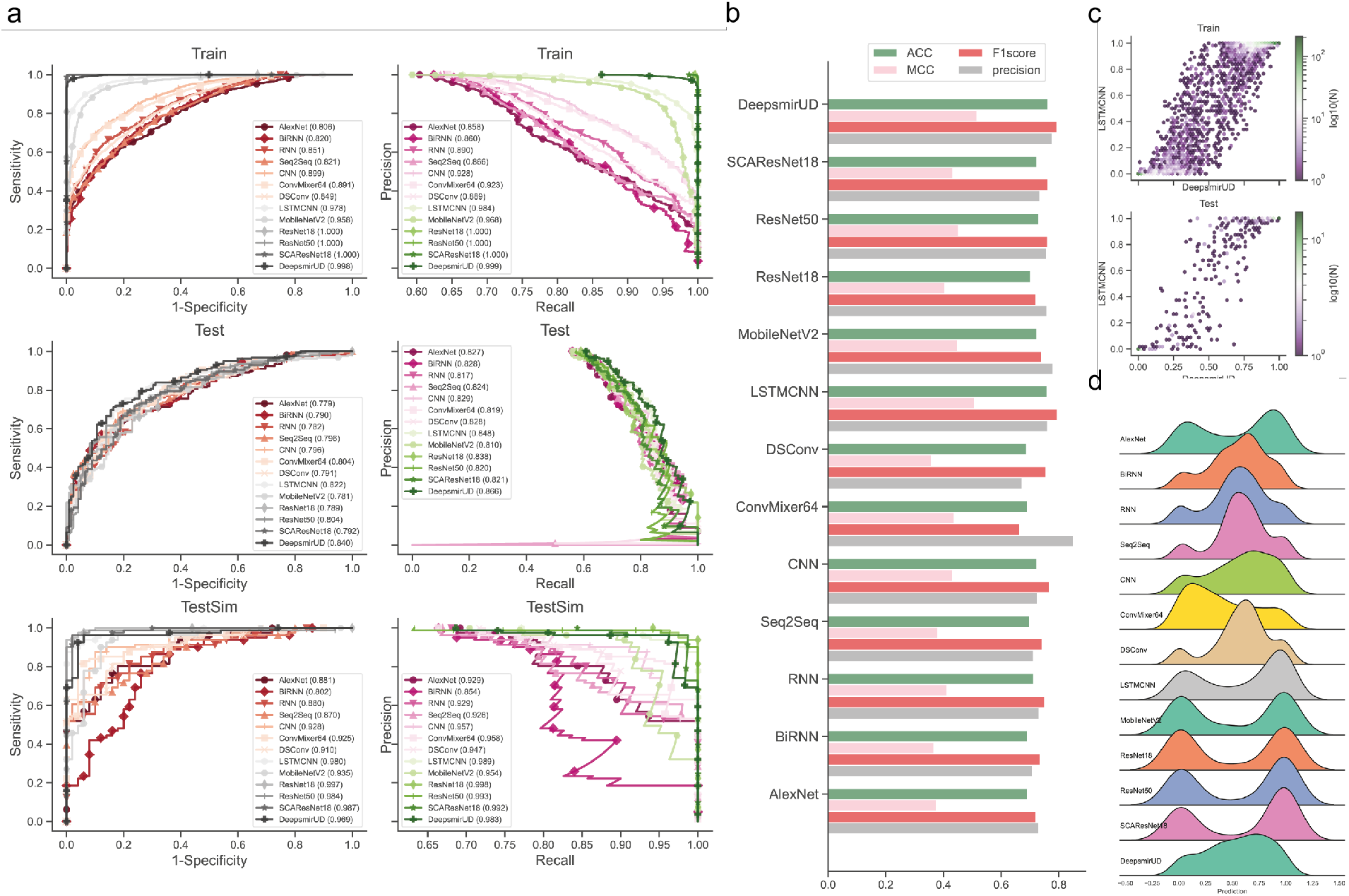
Performance evaluation of the resulting deep learning models on independent test SM-miR relations. a. ROC and precision-recall (PR) curves on the Train, Test, and TestSim datasets. b. ACC, MCC, F1 score, and precision on Test. c. Hexagonal binned plot of comparison between predictions of two leading models (LSTMCNN and DeepsmirUD) on Test. d. Ridge plot of prediction values (i.e., regulatory effects) on Test.

### Regulatory effect prediction on guilt-by-association-formed SM-miR relations

Built outside of experimentally verified relations, the potential SM-miR relations with high confidence are more likely to be unpredictable due to usually a full drug-target relation space in fathomless size. To confer an ability of predicting such potential upregulated and downregulated relations, we further trained our models on samples mixed with SM-miR relations formed using guilt by association (introduced in section *Methods*) and tested them on the TestSim dataset. Surprisingly, all of 13 models achieve extremely high predictive performance with the largest AUC and AUCPR values of up to 1 appearing on most of ResNet-wise methods (**Figure 4a, Table 2, and Supplementary Figures S17b and S18**). BiRNN performs significantly worse than any other methods. This suggests that most of our deep learning algorithms could be prone to well capture the characterization and difference of upregulated and downregulated SM-miR relations in a fine-grained manner.

**Table 2.** Prediction performance evaluation on TestSim.

| Method | AUC | AUCPR | ACC | bACC | Precision | Recall | MCC | F1score | Jaccard |
| --- | --- | --- | --- | --- | --- | --- | --- | --- | --- |
| AlexNet | 0.881 | 0.929 | 0.786 | 0.758 | 0.798 | 0.877 | 0.538 | 0.835 | 0.717 |
| BiRNN | 0.802 | 0.853 | 0.779 | 0.729 | 0.760 | 0.938 | 0.524 | 0.840 | 0.724 |
| RNN | 0.880 | 0.929 | 0.809 | 0.781 | 0.811 | 0.901 | 0.588 | 0.854 | 0.745 |
| Seq2Seq | 0.870 | 0.926 | 0.756 | 0.699 | 0.738 | 0.938 | 0.472 | 0.826 | 0.704 |
| CNN | 0.928 | 0.957 | 0.817 | 0.787 | 0.813 | 0.914 | 0.605 | 0.860 | 0.755 |
| ConvMixer64 | 0.925 | 0.958 | 0.863 | 0.874 | 0.944 | 0.827 | 0.729 | 0.882 | 0.788 |
| DSCnv | 0.910 | 0.947 | 0.794 | 0.753 | 0.781 | 0.926 | 0.555 | 0.847 | 0.735 |
| LSTMCNN | 0.980 | 0.989 | 0.924 | 0.919 | 0.938 | 0.938 | 0.838 | 0.938 | 0.884 |
| MobileNetV2 | 0.935 | 0.954 | 0.893 | 0.887 | 0.914 | 0.914 | 0.774 | 0.914 | 0.841 |
| ResNet18 | 0.997 | 0.998 | 0.969 | 0.968 | 0.975 | 0.975 | 0.935 | 0.975 | 0.952 |
| ResNet50 | 0.984 | 0.993 | 0.931 | 0.933 | 0.962 | 0.926 | 0.857 | 0.943 | 0.893 |
| SCAResNet18 | 0.987 | 0.992 | 0.947 | 0.941 | 0.951 | 0.963 | 0.886 | 0.957 | 0.918 |
| DeepsmirUD | 0.969 | 0.983 | 0.954 | 0.951 | 0.963 | 0.963 | 0.903 | 0.963 | 0.929 |

### Regulatory effect examination using recurrent miRNAs or small molecule compounds

Next, we turned to examine the ability of our methods in predicting regulatory effects of SM-miR relations where either their miRNAs or SMs appear at least once in training SM-miR relations. Using TestRptSM, the top four best-performing methods, DeepsmirUD, LSTMCNN, DSConv, and ConvMixer64 achieve the largest AUC and AUCPR values in the vicinity of 0.770 (**Figure 5a and Tables 3 and 4**). Using TestRptMir, DeepsmirUD and LSTMCNN remain the best-performing methods, with AUC values of 0.909 and 0.918 and AUCPR values of 0.917 and 0.894. Overall, our deep learning algorithms have given good performance using the recurrent miRNAs or small molecules. This further implies that DeepsmirUD is able to be practically applied in screening and discovering drugs targeting a given miRNA and its expression type on a large scale, considering that the training dataset includes many widely-used small molecules and miRNAs.

**Table 3.** Prediction performance evaluation on TestRptMIR.

| Method | AUC | AUCPR | ACC | bACC | Precision | Recall | MCC | F1score | Jaccard |
| --- | --- | --- | --- | --- | --- | --- | --- | --- | --- |
| AlexNet | 0.864 | 0.872 | 0.745 | 0.747 | 0.697 | 0.852 | 0.505 | 0.767 | 0.622 |
| BiRNN | 0.821 | 0.848 | 0.691 | 0.694 | 0.639 | 0.852 | 0.407 | 0.730 | 0.575 |
| RNN | 0.868 | 0.887 | 0.691 | 0.692 | 0.656 | 0.778 | 0.390 | 0.712 | 0.553 |
| Seq2Seq | 0.888 | 0.901 | 0.764 | 0.767 | 0.684 | 0.963 | 0.578 | 0.800 | 0.667 |
| CNN | 0.878 | 0.885 | 0.745 | 0.747 | 0.710 | 0.815 | 0.497 | 0.759 | 0.611 |
| ConvMixer64 | 0.861 | 0.877 | 0.727 | 0.726 | 0.773 | 0.630 | 0.460 | 0.694 | 0.531 |
| DSCnv | 0.832 | 0.866 | 0.691 | 0.695 | 0.625 | 0.926 | 0.438 | 0.746 | 0.595 |
| LSTMCNN | 0.918 | 0.894 | 0.855 | 0.855 | 0.828 | 0.889 | 0.711 | 0.857 | 0.750 |
| MobileNetV2 | 0.888 | 0.891 | 0.782 | 0.782 | 0.778 | 0.778 | 0.563 | 0.778 | 0.636 |
| ResNet18 | 0.836 | 0.836 | 0.745 | 0.745 | 0.760 | 0.704 | 0.491 | 0.731 | 0.576 |
| ResNet50 | 0.807 | 0.813 | 0.673 | 0.673 | 0.655 | 0.704 | 0.347 | 0.679 | 0.514 |
| SCAResNet18 | 0.869 | 0.850 | 0.818 | 0.819 | 0.774 | 0.889 | 0.644 | 0.828 | 0.706 |
| DeepsmirUD | 0.909 | 0.917 | 0.764 | 0.765 | 0.733 | 0.815 | 0.531 | 0.772 | 0.629 |

**Table 4.**
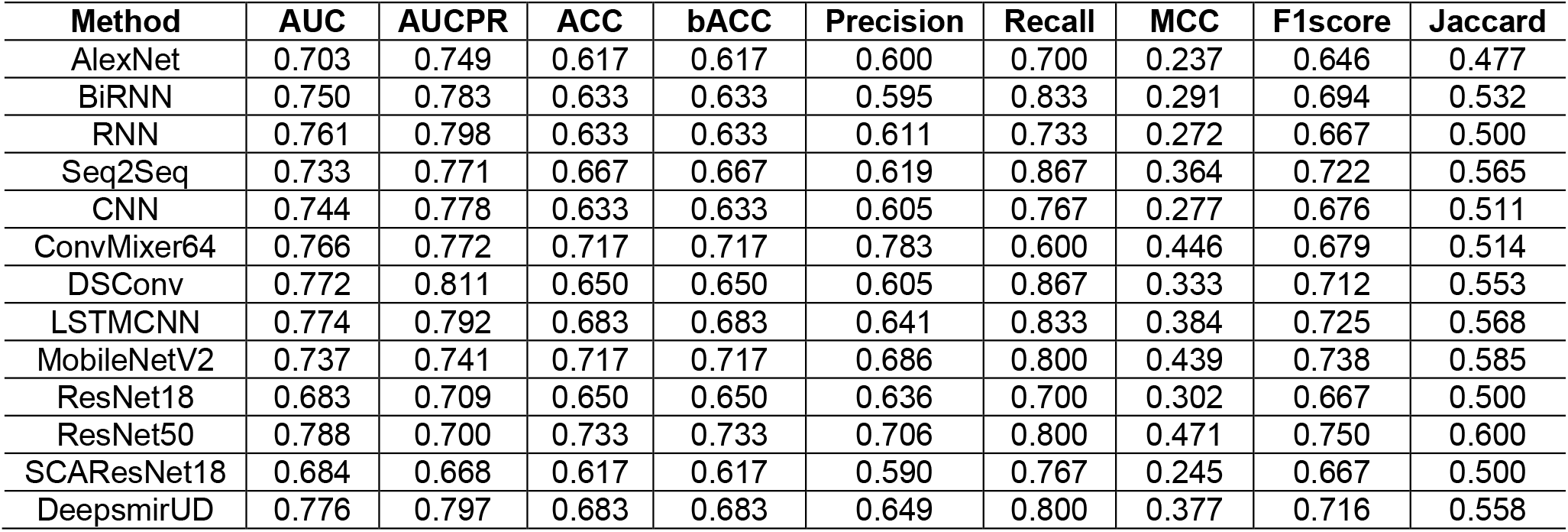
Prediction performance evaluation on TestRptSM.

| Method | AUC | AUCPR | ACC | bACC | Precision | Recall | MCC | F1score | Jaccard |
| --- | --- | --- | --- | --- | --- | --- | --- | --- | --- |
| AlexNet | 0.703 | 0.749 | 0.617 | 0.617 | 0.600 | 0.700 | 0.237 | 0.646 | 0.477 |
| BiRNN | 0.750 | 0.783 | 0.633 | 0.633 | 0.595 | 0.833 | 0.291 | 0.694 | 0.532 |
| RNN | 0.761 | 0.798 | 0.633 | 0.633 | 0.611 | 0.733 | 0.272 | 0.667 | 0.500 |
| Seq2Seq | 0.733 | 0.771 | 0.667 | 0.667 | 0.619 | 0.867 | 0.364 | 0.722 | 0.565 |
| CNN | 0.744 | 0.778 | 0.633 | 0.633 | 0.605 | 0.767 | 0.277 | 0.676 | 0.511 |
| ConvMixer64 | 0.766 | 0.772 | 0.717 | 0.717 | 0.783 | 0.600 | 0.446 | 0.679 | 0.514 |
| DSCnv | 0.772 | 0.811 | 0.650 | 0.650 | 0.605 | 0.867 | 0.333 | 0.712 | 0.553 |
| LSTMCNN | 0.774 | 0.792 | 0.683 | 0.683 | 0.641 | 0.833 | 0.384 | 0.725 | 0.568 |
| MobileNetV2 | 0.737 | 0.741 | 0.717 | 0.717 | 0.686 | 0.800 | 0.439 | 0.738 | 0.585 |
| ResNet18 | 0.683 | 0.709 | 0.650 | 0.650 | 0.636 | 0.700 | 0.302 | 0.667 | 0.500 |
| ResNet50 | 0.788 | 0.700 | 0.733 | 0.733 | 0.706 | 0.800 | 0.471 | 0.750 | 0.600 |
| SCAResNet18 | 0.684 | 0.668 | 0.617 | 0.617 | 0.590 | 0.767 | 0.245 | 0.667 | 0.500 |
| DeepsmirUD | 0.776 | 0.797 | 0.683 | 0.683 | 0.649 | 0.800 | 0.377 | 0.716 | 0.558 |

**Figure 5.**
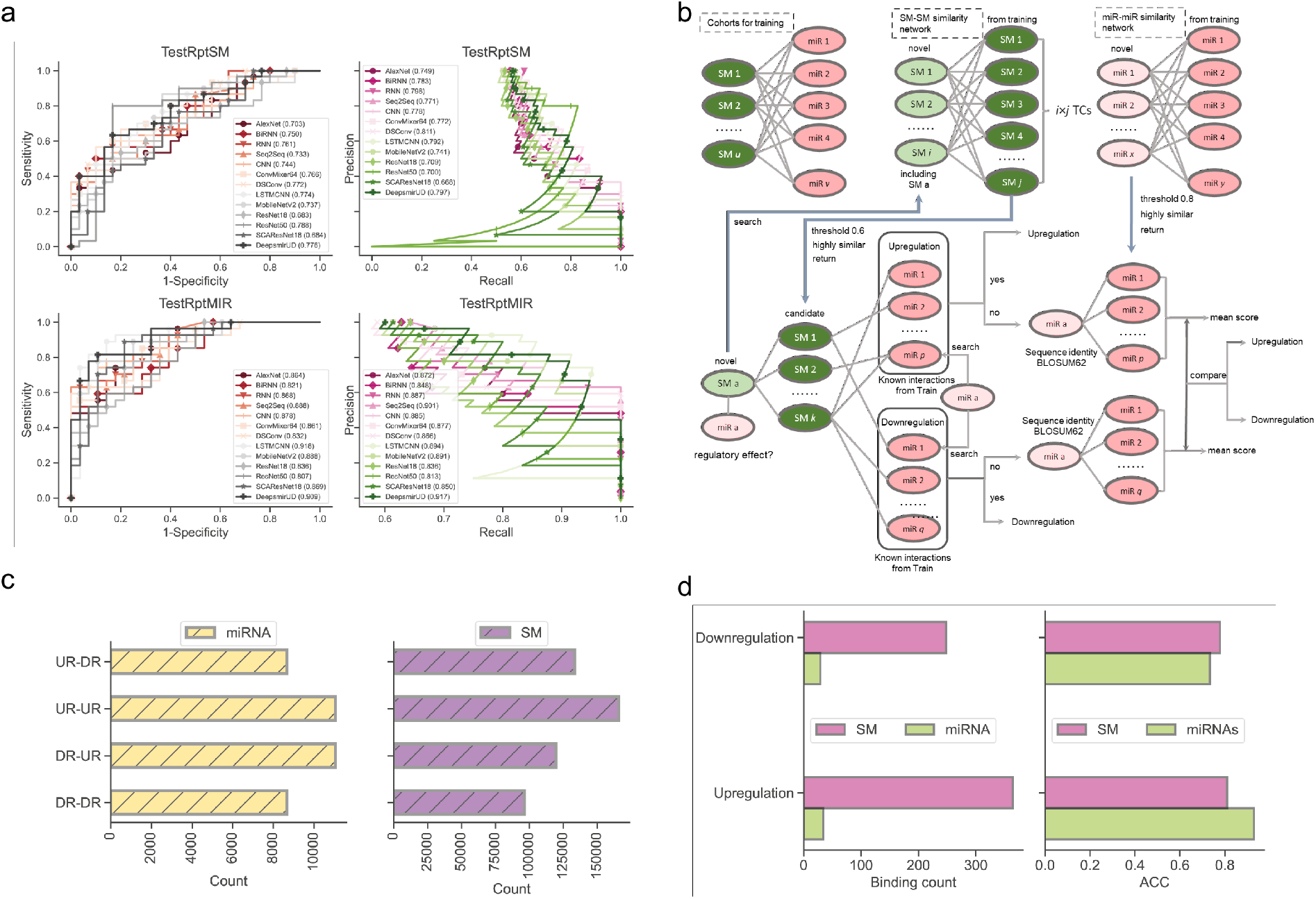
Performance evaluation on the TestRptSM, TestRptMIR, TestUniqSM, and TestUniqMIR datasets. a. ROC and PR curves of deep learning algorithms on the TestRptSM and TestRptMIR datasets. b. Similarity-based network inference of regulatory effects of SM-miR relations formed by novel small molecules or miRNAs. c. Counts of combinations of upregulated novel miRNAs/SMs and downregulated training miRNAs/SMs, upregulated novel miRNAs/SMs and upregulated training miRNAs/SMs, downregulated novel miRNAs/SMs and training downregulated miRNAs/SMs, downregulated novel miRNAs/SMs and upregulated training miRNAs/SMs. The counts reflect the capacity of the two kinds of similarity networks in b (the upper). d. Bar plots of downregulated and upregulated binding counts and ACC.

### Regulatory effect inference using novel miRNAs or small molecule compounds

We then explored how our models were used to predict regulatory effects of SM-miR relations with either novel miRNAs or small molecules. By testing our methods on the TestUniqSM and TestUniqMIR datasets (data not shown), we gained an understanding that these methods were undoubtedly more susceptible to the absence of small molecules in the training data library by offering feeble performance compared to the absence of miRNAs. To exclude a bias made by the random selection of these samples, based on our few trials on altering random seeds 2-3 times, the results were, however, still subject to the above observations. Overall, almost all methods did not actually obtain an ability in precisely predicting the regulatory effect when small molecule compounds are novel, but the 13 models have better AUC values of around 0.5 when miRNAs are novel.

To offer high-confidence inference of regulatory effects, we furthermore constructed a network inference approach based on similarity to assist the optimization-based learning algorithms (see section *Methods*) (**Figure 5b**). Using the network inference approach and relations formed with novel miRNAs, we obtained high ACC values of 0.929 and 0.733 in regard to predicting upregulated and downregulated relations, respectively, while ACC values of 0.810 and 0.778 if using relations formed with novel small molecules (0.813 on average, **Figure 5d**). It should be noted that this network approach does not always return the inference results (i.e., upregulated or downregulated), considering that the inference can be aborted at any steps using the miRNA-miRNA, SM-SM, and/or SM-miRNA networks when no or not enough SMs/miRNAs in the networks are similar to the input SMs/miRNAs.

### Reference map of relations based on gene-expression-perturbed miRNAs or small molecules

To narrow down the gaps in the number of relations between SMs and miRNAs in unexplored space and by experimental verification, the Psmir database employed an enrichment-score (ES) technique to jumps-start not only the inference of SM-miR relations but also the measurement of SM-miR responses (similar to the regulatory effects that we refer to it as in our work). We applied our deep learning methods to infer the upregulated and downregulated associations for SM-miR relations of being in contact, in an effort to expand the knowledge of the unexplored SM-miR relations (**Supplementary Tables 2 and 3**). After stripping out the Psmir-curated associations with a p-value above 0.01 in order to obtain a set of very high-confidence SM-miR associations, we exploited alluvial diagrams to reveal how likely our predicted regulatory effects on miRNA expression with respect to FDA-unapproved and FDA-approved small molecules are similar with the profiles of drug responses to miRNAs in Psmir. Apparently, it can be observed that using a stringent p-value of 0.01 to screen high-quality Psmir relations, the upregulated Psmir relations (FDA-unapproved, overlapped with Train; FDA-unapproved, non-overlapped with Train; FDA-approved, overlapped with Train) largely agree to upregulated relations predicted by DeepsmirUD, while there is the other way around for the downregulated Psmir relations (**Figure 6a and 6d**). The regulatory effects of the SM-miR associations provided in Psmir are predicted as SM-mediated upregulating for miRNAs by most of the deep learning methods (**Figure 6e**, two best-performing methods in **Figure 6c**). For FDA-approved drugs and a p-value of 0.01, we obtained 36 relations predicted as upregulating with 34 unique SMs and 3 miRNAs (non-overlapped), 45 relations predicted as downregulating with 39 unique SMs and 3 miRNAs (non-overlapped), 414 relations predicted as upregulating with 281 unique SMs and 20 miRNAs (overlapped), and 167 relations predicted as downregulating with 148 unique SMs and 20 miRNAs (overlapped). For FDA-unapproved drugs and a p-value of 0.01, we obtained 41 relations predicted as upregulating with 38 unique SMs and 3 miRNAs (non-overlapped), 42 relations predicted as downregulating with 38 unique SMs and 3 miRNAs (non-overlapped), 443 relations predicted as upregulating with 276 unique SMs and 20 miRNAs (overlapped), and 167 relations predicted as downregulating with 141 unique SMs and 20 miRNAs (overlapped). The detailed results can be found in **Supplementary Tables 4-11**. For a visualization purpose, we show our predicted SM-miR downregulation (overlapped with miRNAs in Train, left **Figure 6f**) and upregulation (non-overlapped with miRNAs in Train, right **Figure 6f**) relations formed using FDA-approved small molecules.

**Figure 6.**
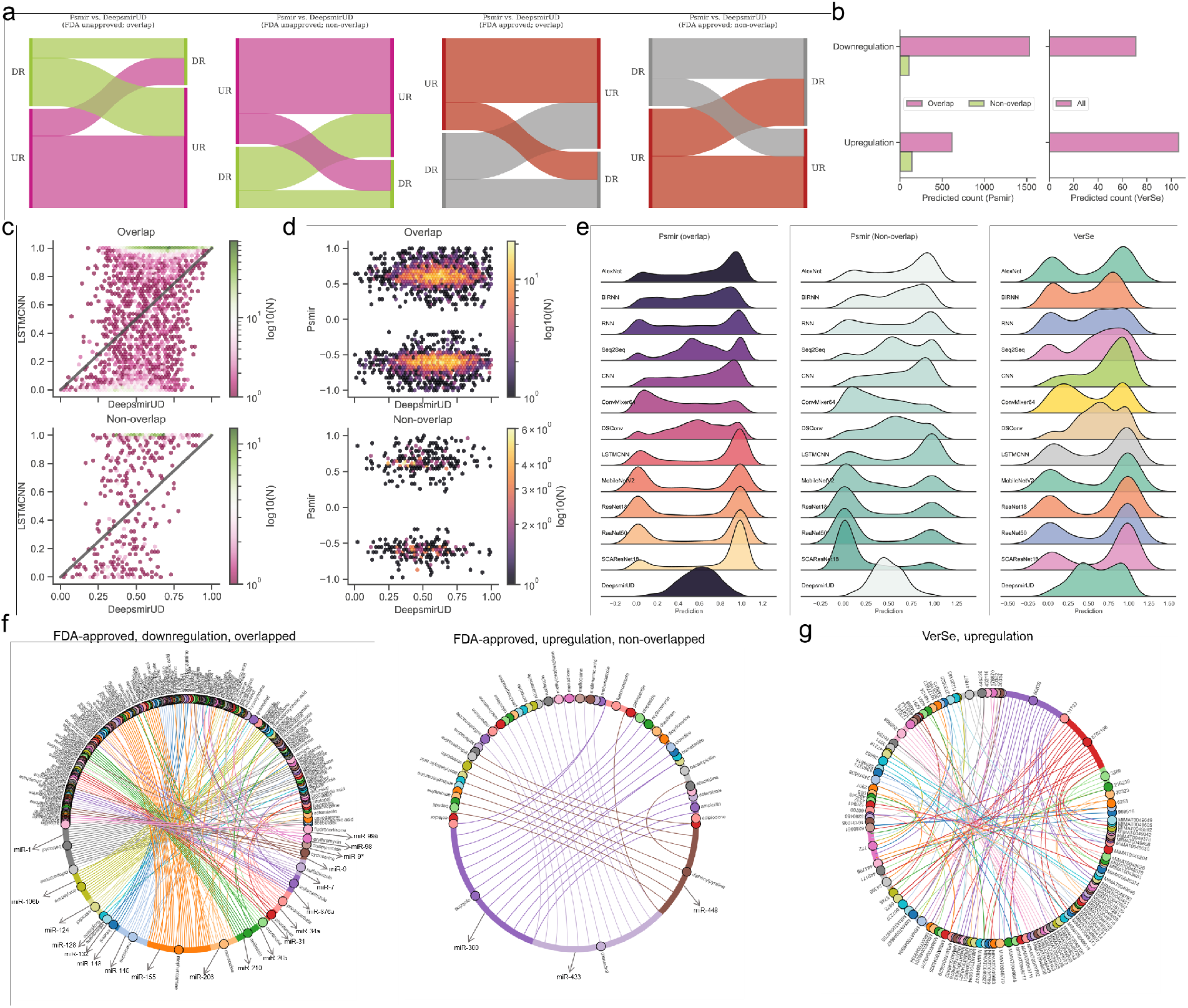
Reference map construction of regulatory effects for SM-miR relations from Psmir and VerSe. a. Alluvial diagram of predicted regulatory effects for FDA-unapproved and FDA-approved relations with miRNAs/SMs that are overlapped and non-overlapped with miRNAs/SMs in Train. b. Prediction of upregulated and downregulated SM-miR relations from Psmir and VerSe using DeepsmirUD. Hexagonal binned plots show a comparison of predicted regulatory effects between DeepsmirUD and LSTMCNN (c), and DeepsmirUD and Psmir (d). e. Ridge plots of predicted regulatory effects. f. Chord diagram of predicted downregulation relations (overlapped with miRNAs in Train, left) and upregulation relations (non-overlapped with miRNAs in Train, right) for Psmir. g. Chord diagram of predicted upregulated relations for VerSe.

### Prediction of regulatory effects of indirect SM-miR relations

Based on 224 curated indirect SM-miR relations screened using miRNA pharmacogenomic data, we further employed our deep learning niche to examine the potential SM-mediated regulatory effects of miRNAs (**Supplementary Table 12**). The approximate 2/3 SM-miR relations are predicted as upregulating concordantly by most of our deep learning methods (**Figure 6b** and **6e**), with 106 upregulated relations versus 71 downregulated relations. The predicted SM-miR upregulation relations are visualized in (**Figure 6g**) and all predictions are made available on DeepsmirUD-web for further use.

### Prediction of SM-disease associations

Potential therapeutic small molecules to treat a cancer disease are able to rectify abnormalities brought about by perturbed profiles of miRNAs in cancer cells, which provides insight into the drug mechanism of action (*i*.*e*., modulating the miRNA expression). By utilizing miRNA-cancer associations from miRCancer and SM-miR relations predicted by DeepsmirUD, we generated SM–disease relationships for drug discovery and repositioning [61–63] to demonstrate the ability of SMs to affect miRNA perturbation profiles in cancers. To evaluate the extent to which a SM can treat a cancer disease, we calculated connectivity scores based on similarity between SM-mediated miRNA perturbation profiles and disease-associated miRNA perturbation profiles (**Figure 6b**, see *Methods* for details). The final predicted associations involve a total of 107 cancers and 1343 small molecules (**Figure 6b**). The resulting negative score suggests the pharmaceutical potential of a SM to treat a disease, while the positive score suggests a cognate perturbation profile between the SM and the disease. Therefore, we retained the SMs with only negative scores and constructed a database to store them at https://rujinlong.github.io/deepsmirud/.

After ranking and extracting high-quality SM-cancer associations with lowest negative connectivity scores, we present two case studies in which the identified druglike small molecules have tangible anti-cancer effects validated by other studies. The FDA-approved antibiotic sulfafurazole has been found to inhibit the tumor growth and metastasis in breast cancer by targeting the endothelin receptor A [64]. Meticrane has been verified to suppress squamous cell carcinoma [65]. Furthermore, we find that many top-ranked associations of single small molecules targeting multiple cancers are predicted to be well-consistent with previous studies. For example, cervical cancer, lung cancer, and colon cancer can be inhibited by naringin [66].

## Materials and methods

### Experienmentally verified SM-miR relations

Similar to existing studies, we derived the SM-miR relations from the SM2miR database (version: April 27, 2015) [27]. After filtering out corrupted data (e.g., entries with out-of-context conditions and repeated entries although found from different publications) from 4989 entries in SM2miR, we first obtained 3641 SM-miR relations, with 2067 upregulated and 1574 downregulated ones. Next, we removed those relations with miRNAs of no FASTA sequences and/or small structures of no SMILE strings. To retrieve FASTA sequences for miRNAs we downloaded the miRbase database (version 22.1) [67] and to fetch SMILE strings from PubChem [68] we used compound identities (CIDs) of small molecules using PubChemPy (https://github.com/mcs07/PubChemPy). All small molecules were confirmed to have certain SMILE strings with a total of 173 unique CIDs presented in the upregulated relation set and 153 unique CIDs presented in the downregulated relation set. After a map from our collected miRNA IDs to the miRbase IDs, we were left with 2037 upregulated relations with 1104 unique miRNAs of known sequences, and 1555 downregulated relations with 867 unique miRNAs of known sequences.

### SM-miR relations based on similarity inference

The full set of SM-miR relations by considering all unique small molecules and miRNAs in SM2miR consists of both experimentally verified relations and unknown relations. If fully exploiting the unknown relations, we might be allowed to identify more of those with great potential to be upregulation and downregulation interacting, to further increase the size of samples for deep learning, and possibly to enhance the generalization abilities of intelligent models due to a diverse composition in training samples. A popular approach to allow for inferring potential binary relations of interest is guilt by association [69,70], which deduces upregulated and downregulated relations based on similarities of drugs and targets.

Similarities between any two miRNAs were measured by sequence identities using the *Pairwise2* module in Biopython (https://biopython.org/) [71] based on the BLOSUM62 matrices of two sequences (detailed in [72]). Due to a restriction of the *Pairwise2* module to be applied only for DNA sequences, we replaced uracil in RNA sequences with thymine ahead of computing identities of any RNA sequence pairs. Similarities between two small molecules were measured using Tanimoto coefficients [73] of their Morgan fingerprints calculated by RDKit (https://www.rdkit.org/) [74]. Any two small molecules are identified similar if the Tanimoto coefficient of the two small molecules is greater than 0.6 as used in [73] and dissimilar, otherwise. Given that miRNA sequences are generally around 20 bases long, we loosened the threshold for judging similarity and dissimilarity between two sequences. Any two miRNAs are identified similar if the sequence identity of the two sequences is above 0.8 (as further discussed in section *Result*) and dissimilar, otherwise.

Inference of whether small molecules and miRNAs interact is performed by guilt by association in a way that interacting/non-interacting SM-miR relations are roughly screened from unknown relations of paired SM-miRs with low/high similarities in comparison to paired SM-miRs in experimentally verified relations. In this process the guilt-by-association approach is applied only once to be able to obtain potentially interacting or non-interacting SM-miR relations [73]. Different from such inference, potentially upregulated and downregulated relations would probably be screened from unknown relations containing three relationships 1) non-interacting, 2) upregulation interacting, and 3) downregulation interacting. To bypass selections that could be erroneously made from non-interacting relations, we screened the potential upregulation and downregulation relations separately by applying the guilt-by-association approach twice for screening high-quality upregulated and downregulated relations, respectively, as shown in **Figure 1a**. Full-connected networks composed of sequence identities between unique miRNAs derived from experimentally verified upregulation and downregulation relations were generated separately (i.e., with guilt by association two times), so were those of unique small molecules. In total, 608,856 miRNA-miRNA sequence identities and 14,878 SM-SM Tanimoto coefficients were calculated to infer potentially upregulated relations while 375,411 miRNA-miRNA sequence identities and 11,628 SM-SM Tanimoto coefficients were used to infer potentially downregulated relations. Consequently, using guilt by association based on a Tanimoto coefficient threshold of 0.6 and a sequence identity threshold of 0.8, we generated 405 and 250 potentially upregulated and downregulated relations, respectively.

### Dataset

To comprehensively investigate the performance of our deep-learning-powered tools, we prepared scores of test datasets stringently (**Figure 1b**). To understand how our tools gain a generalization ability in regulatory effect inference, from the 2037 upregulated relations we first randomly extracted 34 and 368 relations containing 20 unique miRNAs and 20 unique small molecules, respectively. Similarly, we obtained 29 and 252 relations from 1555 downregulated relations. The 34 and 29 relations were combined to be in the TestUniqMIR set and the 368 and 252 relations were finally present in the TestUniqSM set. To test the performance of how sufficiently deep learning approaches were trained, from 1635 remaining upregulated relations we next randomly collected 30 relations (set A) with miRNAs appearing in the remaining relations at least once as well as 30 relations (set B) with small molecules appearing in the remaining relations at least once. The same procedures were repeated to perform another miRNA-orientated pick of 30 relations (set C) and a SM-orientated pick of 30 relations (set D) from 1274 remaining downregulated relations. Set A and set C are combined as TestRptMIR and Set B and set D are combined as TestRptSM. In a ratio of 9:1 we randomly split the left 1575 upregulated and 1214 downregulated relations into upregulated and downregulated ones for training and testing.

In order to enlarge the scale of training data and further augment data, we curated SM-Mir relations from a space of unknown relations criss-crossed by all SMs and miRNAs appearing in the datasets above. The guilt-by-association metric has been introduced to infer potential upregulation and downregulation relations with high confidence from a space of unknown relations. To avoid a lopsided ratio between upregulation and downregulation training samples, the aforementioned 405 and 250 potentially upregulated and downregulated relations were randomly split by a ratio 4:1 into training and test samples. Taken together, we generated a total of 3033 (1471+1092+324+200) training samples, named the Train dataset and 411 (158+122+81+50) test samples, named the Test dataset. To ensure no overlaps between training and test samples owing to the addition of guilt-by-association relations for training, we finally detected and removed 2, 5, and 7 overlapped relations from Test, TestRptMIR, and TestUniqSM, respectively. Detailed information about all datasets of training and test samples can be found in **Supplementary Table S13-S20**.

### Definition of small molecule-mediated regulatory effects on miRNA expression

In this work, the small molecule-mediated regulatory effect on miRNA expression refers to the likelihood of a downregulation or upregulation SM-miR relation pair, *i*.*e*., how likely the expression of a miRNA is downregulated or upregulated by a small molecule. Deep learning models are optimized and set to learn the characterization of downregulation or upregulation relation pairs in order to output their likelihoods as much accurately as possible.

### Feature representation

Our deep learning approaches make use of a 1369-size feature vector to characterize the relation between a small molecule and a miRNA (**Figure 2**). It reflects the positional, compositional, physicochemical, or structural properties of the small molecule or the miRNA, with the former represented by a vector of length 223 and the latter represented by a vector of length 37. Given that square-shaped filters are more popularized in computer vision to perform feature extraction on an input object with a dimension of usually greater than one, we then converted each feature vector to an image-like matrix with a 37×37 dimension and an initial channel equal to one for computer-vision methods including AlexNet, CNN, ConvMixer, DSConv, LSTMCNN, MobileNetV2, ResNet, and SCAResNet18, while at a single time step we sequentially cropped out a 37 input size from the feature vector, with a total of 37 time steps for speech-recognition methods including RNN, BiRNN, and Seq2Seq. The features of small molecules and miRNAs are listed below.

The miRNA vector is comprised of positional and compositional features.

#### *Nucleotide Composition* (NAC)

NAC measures the percentage of a single nucleic acid type in a nucleotide sequence, which is computed by

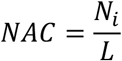

where *L* represents the length of a miRNA sequence and *N*_*i*_ represents the total count of a nucleotide *i* in the sequence.

#### *Di-Nucleotide Composition* (DNC)

DNC measures the percentage of the combination of any two nucleotides in a nucleotide sequence, given by

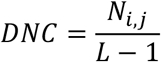

where *N*_*i,j*_ represents the total count of a pair of nucleotides *i* and *j* in the sequence.

#### *Tri-Nucleotide Composition* (TNC)

TNC is used to describe the composition of 64 unique triplets of nucleotides for a sequence, which is expressed as

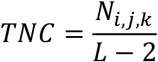

where *N*_*i,j,k*_ represents the total count of a triplet of nucleotides *i, j* and *k* in the sequence.

#### *Qua-Nucleotide Composition* (QNC)

TNC is used to describe the composition of 256 unique quadruplets of nucleotides for a sequence, which is expressed as

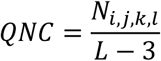

where *N*_*i,j,k,l*_ represents the total count of nucleotides *i, j, k*, and *l* in the sequence.

#### *Composition of t-Spaced Nucleoic Acid Pairs* (CTSNAP)

CTSNAP [75] is a 16-size vector containing the percentages of 16 possible pairs of nucleotides with a *t* distance apart, which are defined as

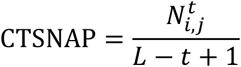

where 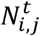 represents the total count of a pair of two nucleotides (*i, j* ∈ AA, AC, AG, AT, CA, CC, CG, CT, GA, GC, GG, GT, TA, TC, TG, TT) in distance *t*.

#### *Average Accumulated Nucleotide Frequency* (aveANF)

We define aveANF, which encodes both positional and compositional information for a given nucleotide *i*, given by

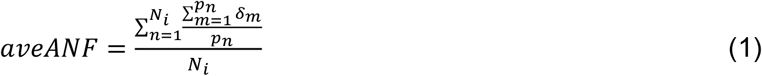

where *p*_*n*_ represents the position of the nucleotide *i* and *δ*_*m*_ is the Kronecker symbol to indicate if a nucleotide *i* is present at position *m*,

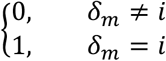

#### Fingerprint

The Morgan fingerprint of length 1024 and the density of the Morgan fingerprint were encoded into vectors for small molecules.

#### Structural and physiochemical encoding

We extracted from each small molecule 6 structural elements, including the number of rings, the number of heavy atoms, the number of nitrogens and oxygens, the number of NHs or OHs, the number of aliphatic carbocycles, and the number of heteroatoms. Besides, the maximum and minimum of Estate Indexes and the Waals surface area combining Estate were considered. Finally, LogP and molar refractivity reflecting compound’s physiochemical properties were also encoded.

All miRNA features are implemented based on Python and all small molecule features are generated using RDKit (https://www.rdkit.org/).

### Performance assessment

Our deep-learning approaches are evaluated by threshold-free measurements including AUC and AUCPR [76] and a number of threshold-based measurements, including accuracy (ACC), balanced accuracy (bACC), precision, recall, the Jaccard index, the Matthews correlation coefficient (MCC), and F1 score.

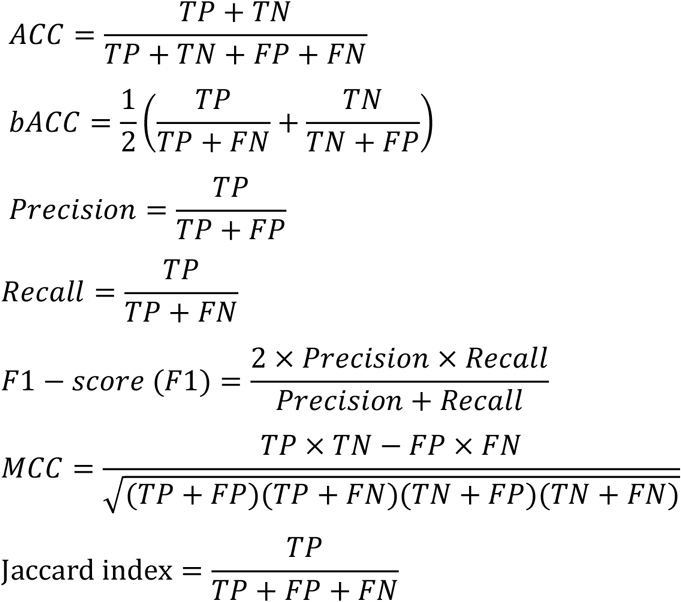

where TP (true positive), FP (false positive), TN (true negative) and FN (false negative) stand for the number of upregulation SM-miR pairs to be predicted correctly as upregulating, the number of downregulation SM-miR pairs to be predicted incorrectly as upregulating, the number of downregulation SM-miR pairs to be predicted correctly as downregulating, and the number of upregulation SM-miR pairs to be predicted incorrectly as downregulating, respectively. For threshold-based methods, a threshold of 0.5 was taken to calculate TP, FP, TN, and FN.

### Deep learning methodology

It is well known that in recent years, the number of off-the-shelf deep learning models has gone into orbit [46], due to not only an increasing amount of input from research communities but also the flexibility in stacking or removing the number of unit neural network structures required for training and optimization as well as in changing other detailed settings in neural layers of note [40]. This is largely different from most of traditional machine learning algorithms relying on relatively fixed model structures. The performance of different deep learning models can also be slightly or largely different, which can be evidenced partially by a plethora of computer-science publications with different models to annually compete for better prediction abilities on standard training cohorts such as CIFAR-10 [47]. Without a detailed benchmark, choices of deep learning models are unknown. For this reason, we opted for 6 existing and well-established deep learning models (Alexnet, ConvMixer, MobileNetV2, ResNet18, ResNet50, and SCAResNet18) and 6 our assembled architectures (BiRNN, CNN, DSConv, LSTMCNN, RNN, and Seq2Seq) and comprehensively evaluated their capability of predicting SM-mediated regulatory effects on miRNA expression in expectation of a better performance gain.

#### Alexnet

Alexnet was first introduced to perform an image recognition task in the LSVRC-2010 contest, hoving widely into sight by achieving the state-of-the-art performance [44]. The backbone of Alexnet consists of 5 convolutional layers and 3 fully-connected layers. We kept using the raw setting of Alexnet, with 96 filters of size 11×11, 256 filters of size 5×5, 384 filters of size 3×3, 384 filters of size 3×3, and 256 filters of size 3×3 placed in the respective convolutional layers in order.

#### BiRNN

The bidirectional recurrent neural network (BiRNN) is a variant version of the recurrent neural network (RNN), which can be trained in forward and backward time directions simultaneously [51]. Our BiRNN structure consists of a single BiRNN layer preceded with two RNN layers, each with the same 256 hidden units.

#### CNN

As one of the most typical deep learning components, convolutional neural networks (CNN) have been pervasively used in multiple kinds of fields [49]. We constructed a 3-layer CNN structure, each followed by a max pooling operation with a step of 2. Finally, a 128-neuron fully-connected layer was placed after completing convolutions. We applied 32 filters of size 3×3 to extract features from input.

#### ConvMixer64

Very recently, the ConvMixer model [46] has been proposed as a type of vision transformers (ViTs) to handle image recognition by learning the patches of image-like objects. The ViT module can remove inductive biases of convolution operations to boost performance to some extent, thereby starting to be popularly applied in computer vision. We kept the backbone of the ConvMixer model the same as in [46] but replaced 256 filters with 64 filters for feature extraction and reduced the number of ConvMixer blocks from 8 to 2, in order to reduce computing power required for training at a manageable level.

#### DSConv

In the Xception method, the recurrent depthwise-separable convolution (DSConv) modules are described as the cornerstone of the high performance of image recognition [52,77]. The key of DSConv lies in a hypothesis that the full-scale separation between cross-channels correlations and spatial correlations in relation to the feature maps of CNN may have positive impact on performance increases. This has been tested efficient to improve image recognition performance. We were interested in whether our study can, to some extent, benefit from the use of the DSConv operation alone and thus constructed a DSConv-based deep learning framework consisting of solely the DSConv operation that alternates between a depthwise convolutional layer and a separable convolutional layer 2 times, which finally connects to a 128-neuron fully-connected layer. The max pooling operation with a size of 2 was performed only after a depthwise convolutional layer. Likewise, 32 filters of size 3×3 were applied to extract features from input data.

#### LSTMCNN

A LSTMCNN layer is a hybird of convolutioanl layer and a long short-term memory (LSTM) layer [50,53]. The output of the LSTM component is a convolution-like transformation that suits the input to the next convolutional layer. We constructed a structure with 3 LSTMCNN layers, each followed by a max pooling operation with a step of 2. We applied 32 filters of size 3×3 to extract features from input. We used the ConvLSTM2D module of Tensorflow (https://www.tensorflow.org/) as each LSTMCNN layer.

#### MobileNetV2

MobileNetV2 was designed for object recognition tasks in mobile and resource-constrained environments and has been shown to effectively remove non-linearities in the layers, demonstrating high predictive performance [45]. We introduced the raw MobileNetV2 model into the regulatory effect inference. In order to maintain the acceptable consumption of CPU and memory resources, on every single convolutional layer we kept the filter number of no greater than 64.

#### ResNet18

Residual neural networks (ResNets) have gained popularity in deep learning applications and have recently achieved great success in protein structural prediction [38]. A well-tried ResNet with 18 residually-connected convolutional layers used in [47] was adopted as our ResNet18 method.

#### ResNet50

Our previously two methods in predicting residue contacts [78] and inteactions sites [76] in transmembrane proteins have benefitted from the use of ResNets with massive layers (>35). Therefore, in addition to the 18-layer ResNet, we also applied a 50-layer ResNet (used in [47]) to in-depth extracting features and learning representations of the SM-mediated regulatory effects on miRNA expression.

#### RNN

The recurrent neural network (RNN) is one of the basic structures of deep learning algorithms, used quite commonly in speech and language processing [50,79]. Our RNN structure begins with 2 LSTM-type RNN layers, each with 256 neurons inside. It finally connects to a fully-connected layer with 256 nerons.

#### Seq2Seq

The sequence-to-sequence method has been introduced in modeling natural language semantically and syntactically [55]. It encodes and decodes a sequence of input symbols (i.e., feature vectors) and has been tested effective to learn the representations of the input symbols [56]. At the core of our Seq2Seq structure are one RNN-like encoder and one RNN-like decoder, followed finally by a fully-connected layer with 256 neurons. The encoder has exactly the same structure of our BiRNN algorithm, so does the decoder but with the 3 layers placed in reverse order.

#### SCAResNet18

The attention mechanism is brought up for recasting and refining intermediate feature maps locally so as to enhance representational power of deep learning algorithms in the spatial and channel-wise directions, leading to two derivatives, spatial and channel-wise attention (SCA) modules introduced by Woo et al. [48]. We adopted one of their reported models, CBAM-ResNet18 integrating ResNet18 with the SCA modules. We term it SCAResNet18 in our study.

### Ensemble of deep learning models

DeepsmirUD —the final ensemble model— was obtained by averaging the predictions produced by the 12 deep learning models on each dataset, which reduced variations among predictions [76].

### Training deep learning algorithms

As akin to our previous work [76,78], we picked the Adam method [80] for optimization of all deep learning algorithms with a learning rate 1e-3 and a batch size of 100. The parameters of the deep learning methods were estimated based on 5-fold cross validation. The categorical cross entropy was introduced to measure how different ground-truth and predicted SM-miR relations were.

### Overfitting prevention

Overfitting [81] has a prominent impact on lowering generalization abilities of intelligent models on unseen SM-miR relations, especially those formed with small molecules and/or miRNAs to be largely distinguishable in features to every single small molecule and miRNA in training samples. To avoid overfitting, we adopted the early stopping strategy (previously used in [76]) to pick out sufficiently trained models in a way that training had no sooner terminated than the prediction performance showing a falling tendancy on test/validation data.

### Network inference approach for novel SM drugs and miRNA targets based on smilarity

We further proposed a network inference approach for accurately predicting regulation types of SM-miR relations formed with either novel small molecules or novel miRNAs. Since the size of cohorts (∼3000) for training was quite limited and the difference between training and unseen SM chemical structures can be very high, the deep learning power might be impaired when either unseen small molecule or unseen miRNA was used. This approach was built by taking advantage of training sample information in the following ways. First, we built two similarity networks, one storing SM-SM similarity scores between all unique small molecules and all training small molecules and the other one storing miRNA-miRNA similarity scores between all unique miRNAs and all training miRNAs (**Figure 5b**). For a predicted or experimentally-verified relation, the network inference approach can select a small molecule or a miRNA as input. As an example, we assumed a query SM-miR relation formed by a novel small molecule and we began by selecting the small molecule as input to search all small molecules from the SM-SM similarity network if any of them shared a pre-defined threshold of 0.6 with the small molecule. Then, relations containing the returned small molecules were picked from both sets of known upregulation and known downregulation relations (from the Train dataset) and had their miRNAs matching the partner miRNA of the query small molecule. If a perfect match appears in the known upregulated/downregulated relations, the query relation was correspondingly assigned upregulation/downregulation. Otherwise, two subnetworks containing similarity scores between the partner miRNA and those picked upregulated and downregulated miRNAs are extracted from the miRNA-miRNA similarity network, respectively. Finally, the similarity scores in each subnetwork are averaged and the bigger one decides whether the query relation is upregulated and downregulated. We repeated the same workflow for relations formed by novel miRNAs but we screened the candidate miRNAs based on a query miRNA from the miRNA-miRNA similarity network with a threshold of 0.8. Because the miRNAs of small sequence length are more amenable to causing a lot of high scores of similarities between miRNAs than the very long fingerprint lengths of small molecules. This approach is also included in the DeepsmirUD toolkit.

### Psmir relations miRNA-perturbed gene expression profiles

Psmir is an archive of high-confidence predicted SM-miR relations selected from a curated collection of SM-miR candidate relations constructed with miRNAs from miRNA-perturbed gene expression profiles in the GEO database and small molecules from small molecule-perturbed gene expression profiles in cmap [82]. With a pool of 51,051 candidate SM-miR relations formed by 1309 unique small molecules and 39 unique miRNAs, 6501 relations containing 1295 unique small molecules and 25 unique miRNAs were screened as interacting by applying a statistically significant p-value <=0.05. We identified 1195 small molecules whose compound names were successfully matched using PubChemPy with certain small strings. We removed those relations sharing the same miRNAs but with different compound names that were converted into the same CIDs, leaving 5001 SM-miR relations. After removal of relations with miRNAs in no match to any records in miRBase (in order to obtain precise miRNA sequences), we finally obtained 4656 SM-miR relations of which 4156 (overlapped) have their miRNAs having appeared in our training relations and 500 (non-overlapped) have unique miRNAs to any miRNA in our training relations. We also examine the high-quality set of relations with a p-value no greater than 0.1. The negative or positive response of drugs to miRNAs in Psmir was determinded by a negative or positve association score (AS), respectively.

### VerSe with pharmacogenomic miRNAs

VerSe supplies the painstaking curation of miRNA pharmacogenomic sets that essentially embody miRNA-gene-drug triplets in a ternary relationship in which miRNAs inhibit genes and further affect the response of drugs that are relevant with the genes [83]. We attempt to dig up potential influence of miRNAs/drugs on drugs/miRNAs from a collection of 272 SM-miR indirect links collected from a total of 140 publications in VerSe. Note that VerSe does not provide whether SM-miR relations are upregulations or downregulations and SMs and miRNAs are indirectly related. After examining available sequences of the miRNAs using miRBase we were left with 257 SM-miR indirect links and after eliminating links with drugs of no smile strings we finally obtained 224 SM-miR indirect links. Similar to the deduplication procedure used in Psmir, after removing relations sharing the same miRNAs but different compound names mapped onto the same CIDs using PubChemPy, we finally retained 177 SM-miR indirect links.

### miRNA-cancer database

Cancer-related miRNA data is obtained from the curated miRCancer database (version: 08.27.2019) [57], which contains 9080 relations in 196 cancer types from 7288 papers. This database includes the upregulated and downregulated miRNAs information for each cancer type.

### Connectivity scoring of SM-cancer associations

Druglike capabilities of SMs in treating cancers are evaluated according to connectivity scores (ranging from -1 to 1) calculated with the Weighted Kolmogorov-Smirnov (WKS) method [84], which is available at https://github.com/Jasonlinchina/RCSM/R/GSEAweight1.R. A SM with a negative score represents its mediated miRNA perturbation signature opposite to that of a query disease and has new therapeutic potential as a drug against the disease, while a positive score suggests the similarity between the SM-mediated and the disease-associated miRNA perturbation signatures [62,85]. As seen in **Figure 7a**, with a given disease we first retrieved all involved genes transcribing cancer-related miRNAs and split them into two sets, one being upregulated and the other one being downregulated in the disease. Then, we built a SM×miRNA matrix based on upregulated and downregulated SM-miR relations predicted by DeepsmirUD. In this matrix, each SM’s miRNA perturbation signature is represented by encoding its relations with all used miRNAs. Finally, the disease miRNA signatures and the ranked miRNA signatures for each SM are compared to yield a series of potential drugs with negative connectivity scores. Note that the SM×miRNA matrix was built by taking advantage of all SM-miR relations used in this study in order to comprehensively screen drugs.

**Figure 7.**
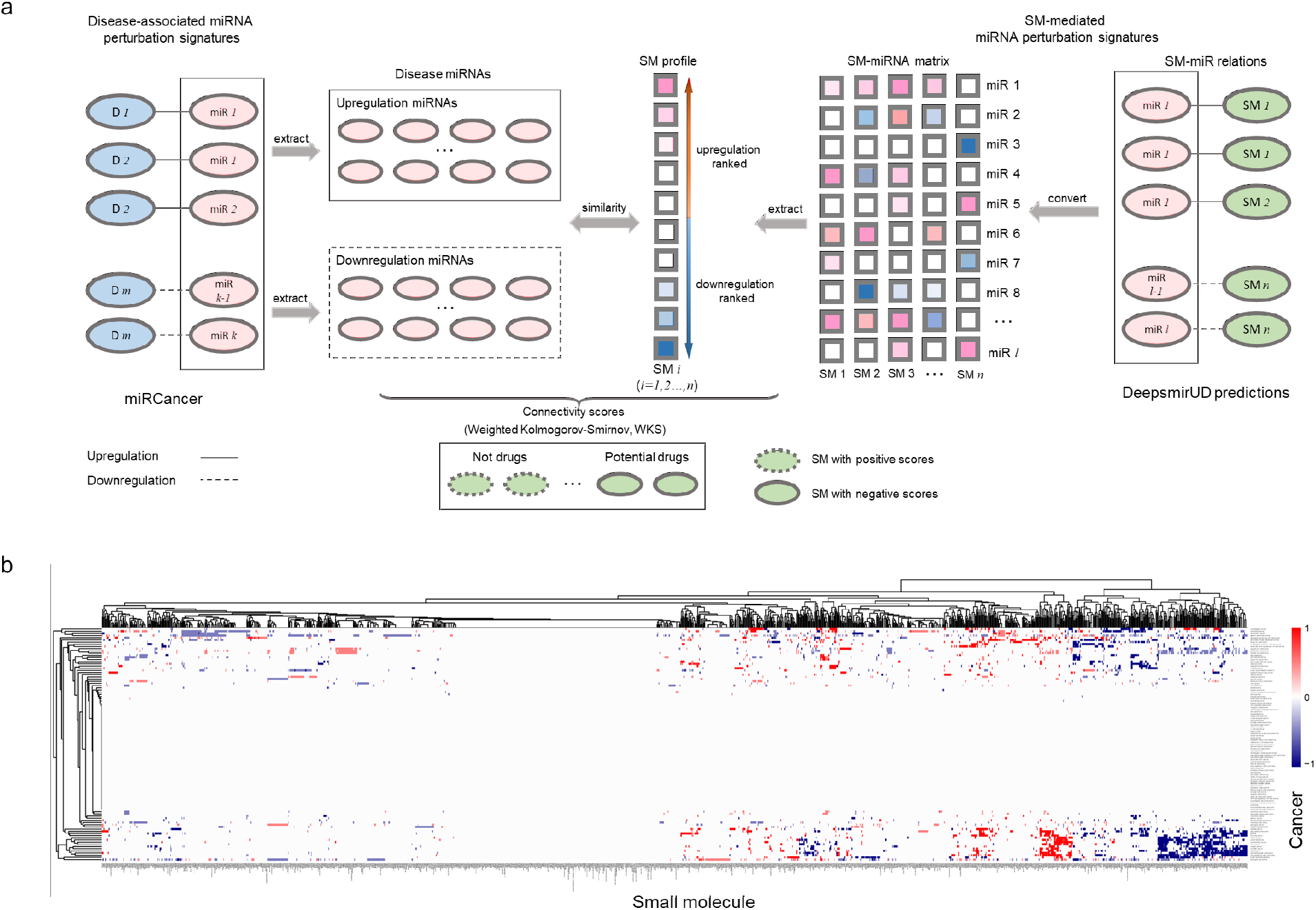
SM-cancer connectivity analysis. a. Illustration of connectivity scoring to infer SM-cancer associations. b. Heatmap of connectivity between 1343 small molecules and 107 cancer types. The positive connectivity score represents similar perturbations between a disease signature and a drug profile, while the negative connectivity score stands for the drug reversal of the disease signature. Note that a well-resolved version of the SM-cancer heatmap can be fetched from https://rujinlong.github.io/deepsmirud/.

## Discussion

Here, we discuss possible explanations about contributing to stable performance by deep learning. The variation of characterizing miRNA sequences (*e*.*g*., using composition) of small length is much more measurable and controllable compared to longer transcriptome sequences. Therefore, the deep learning architectures may better capture and learn representations from miRNA sequences to favour better performance comparatively. We applied a fairly rigorous pre-processing procedure to select high-quality SM-miR relations, which effectively eliminates noisy samples. Existing studies have revealed that overfitting of models is a common and key factor to impair model generalization abilities on unseen data in practical application stages. To eliminate such an influence that occurred during training, we monitored and visualized the entire landscapes of performance variations over 400 epochs. Our final models of choice have excluded those deep learning architectures with their trained models showing a notable and long-term fluctuation over epochs, which was treated as a relatively unstable or weak learning process of features from the SM-miR downregulated and upregulated relations. Mechanistically, due to flexible assembly of layers, there can be a hefty number of different deep learning architectures and their performance can differ slightly or largely. Thus, we managed to collect various models in search of the optimum prediction capability. Overall, due to a high flexibility in deep learning structure settings, it is computationally hard to confirm best combinations of parameters of neural network layers. In a manageable setting of allowing for computing and memory resources, we have attempted to provide high-quality models from many competing and canonical deep learning frameworks to assist the accurate and fast examination of SM-mediated regulation types for miRNA expression. Pharmaceutically, DeepsmirUD can be applied in three scenarios as follows.

1. Biologists have needs of shrinking the search range of small molecules to drug a disease-specific miRNA target for specific regulation types.
2. For a small molecule of interest, upregulation and downregulation types of miRNAs induced by this small molecule can be identified.
3. With given gene family-specific miRNA targets, a pool of their potential small molecule drugs can be examined.

Besides, as small molecules widely target RNAs from non-coding regions and the sequence composition or similarity of ncRNAs of small length may bear resemblance to miRNAs’ than longer ncRNAs’, we estimate that the application of DeepsmirUD could be further extended to predicting the SM-mediated regulatory effects on other types of ncRNAs such as siRNAs.

Disease can be treated by targeting associated miRNAs with small molecules, which can help find novel drug-disease relations. This provides direct biomedical practice for leveraging the upregulation and downregulation relations predicted by DeepsmirUD. We have demonstrated the utilization of miRCancer combined with connectivity scoring to mine the novel drugs. We foresee that our results give novel insight into drug repurposing and discovery by taking advantage of SM-miRNA-disease relationships.

## Conclusion

In this paper, we have presented a novel approach to quantify the small molecule-mediated regulatory effects on miRNA expression by training 12 deep learning architectures as an ensemble, DeepsmirUD. Through multiple rounds of restricted selections on trained models in order to avoid overfitting, the results show that DeepsmirUD can accurately model the relation types (upregulation and downregulation) of a coupled pair of a small molecule and a miRNA. The inferred regulatory effect levels reflect confidence in how strongly miRNAs are upregulated or downregulated by small molecules, which provide potentially therapeutic choices on miRNAs to be targeted for specific oncogenic pathways. DeepsmirUD differs any existing tool by its uniqueness, flexibility, and predictive power, which is the first method used in an ad hoc manner in this research field to allow a directional screening of upregulation or downregulation miRNAs with an additional benefit in small molecule-based drug discovery. We stress a broad application scenario in which DeepsmirUD can be applied for screening either miRNAs targets or small molecule drugs on a large scale computationally, giving a set of either predicted or experimentally-verified SM-miR relations. In conclusion, our methods are expected to speed up the development of therapeutics to treat disease-associated pathways that miRNA targets regulate.

## Code and data availability

The DeepsmirUD program is publicly available at https://github.com/2003100127/deepsmirud. Predictions of regulatory effects on miRNA expression for a large number of SM-miR relations and cancer-related drugs identified by means of SM-miR relation predictions are available at https://rujinlong.github.io/deepsmirud/.

## Author contribution

J.S., X.W., and Z.C. conceived and designed this research. J.S. performed the computational experiment and the deep learning study. J.R. and X.W. performed the disease-associated analysis. J.S., X.W., and Z.C. curated the data. J.S., X.W., Z.C., and S.C. made the investigation. J.S., J.R., and X.W. visualized the data. J.S. and X.W. analysed the results. J.S. and J.R. developed the software. J.R. developed the DeepsmirUD-Web website. J.S. wrote the manuscript. X.W., J.S., F.Q., L.R.M., A.P.C., and L.D. edited and revised the manuscript.

X.W. managed this project. X.W. and L.D. provided supervision and acquired funding.

## Acknowledgement

This study was supported by Chinese Universities Scientific Fund (Z1090222009). FQ is supported by the National Natural Science Foundation of China (Grant No. 32000462) and the Scientific Research Funds of Huaqiao University (Grant No. 22BS114).

