## Supplementary_material for "DeepsmirUD: Precise prediction of regulatory effects on miRNA expression mediated by small molecular compounds using competing deep learning frameworks"

**Table S1**. Prediction performance evaluation on Train.

| Method | AUC | AUCPR | ACC | bACC | Precision | Recall | MCC | F1score | Jaccard |
|---|---|---|---|---|---|---|---|---|---|
| AlexNet | 0.806 | 0.858 | 0.714 | 0.704 | 0.742 | 0.770 | 0.412 | 0.755 | 0.607 |
| BiRNN | 0.820 | 0.860 | 0.732 | 0.711 | 0.726 | 0.857 | 0.446 | 0.786 | 0.647 |
| RNN | 0.851 | 0.890 | 0.758 | 0.743 | 0.762 | 0.840 | 0.499 | 0.799 | 0.665 |
| Seq2Seq | 0.821 | 0.866 | 0.728 | 0.709 | 0.730 | 0.834 | 0.435 | 0.779 | 0.637 |
| CNN | 0.899 | 0.928 | 0.800 | 0.783 | 0.784 | 0.901 | 0.591 | 0.838 | 0.722 |
| ConvMixer64 | 0.891 | 0.923 | 0.760 | 0.783 | 0.934 | 0.626 | 0.576 | 0.749 | 0.599 |
| DSConv | 0.849 | 0.889 | 0.746 | 0.717 | 0.719 | 0.915 | 0.483 | 0.805 | 0.674 |
| LSTMCNN | 0.978 | 0.984 | 0.913 | 0.910 | 0.916 | 0.934 | 0.822 | 0.925 | 0.861 |
| MobileNet | 0.958 | 0.968 | 0.891 | 0.895 | 0.938 | 0.866 | 0.782 | 0.901 | 0.819 |
| ResNet18 | 1.000 | 1.000 | 0.998 | 0.998 | 0.999 | 0.997 | 0.995 | 0.998 | 0.996 |
| ResNet50 | 1.000 | 1.000 | 0.996 | 0.996 | 0.998 | 0.996 | 0.993 | 0.997 | 0.994 |
| SEResNet | 1.000 | 1.000 | 0.997 | 0.997 | 0.997 | 0.998 | 0.994 | 0.997 | 0.995 |
| DeepsmirUD | 0.998 | 0.999 | 0.978 | 0.978 | 0.982 | 0.980 | 0.956 | 0.981 | 0.963 |

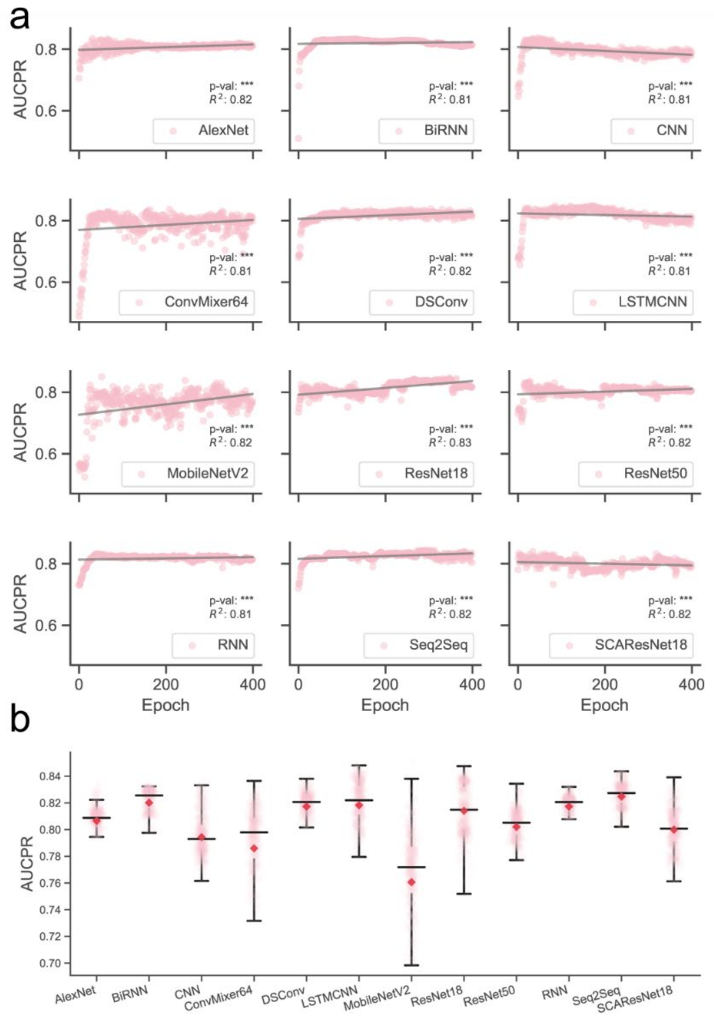


Figure S1. Full-scale AUCPR performance examination of deep learning algorithms on Test over training epochs. a. Landscapes of AUCPR variations. b. Boxplot of AUCPR values.

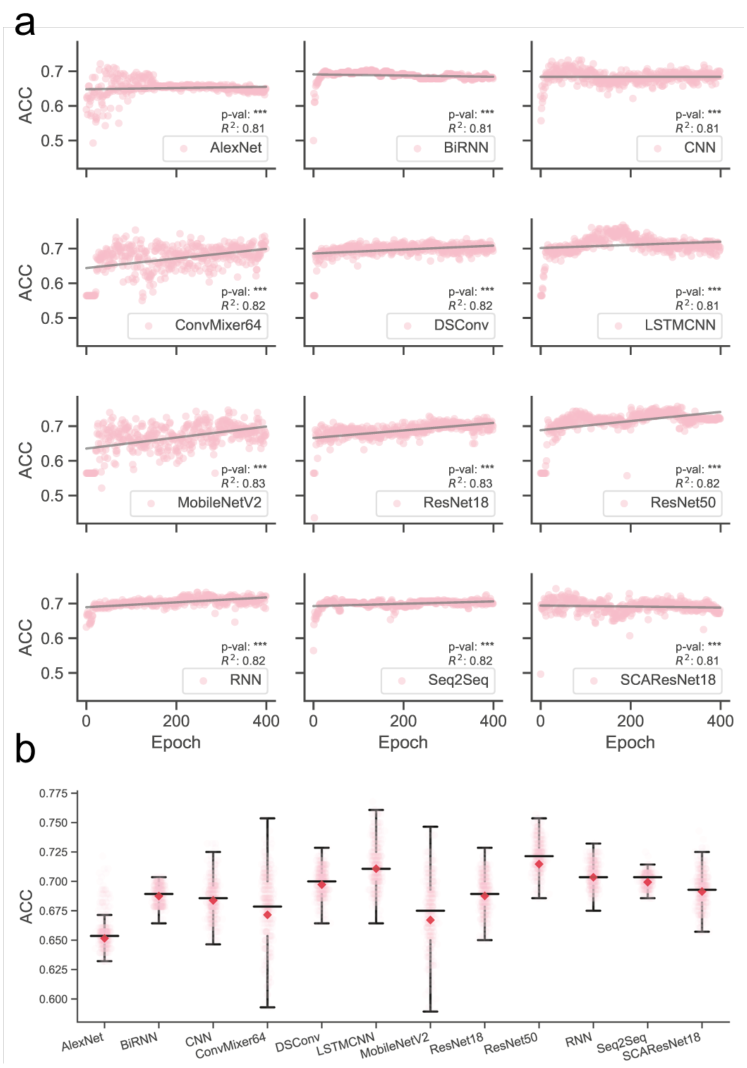


Figure S2. Full-scale ACC performance examination of deep learning algorithms on Test over training epochs. a. Landscapes of ACC variations. b. Boxplot of ACC values.

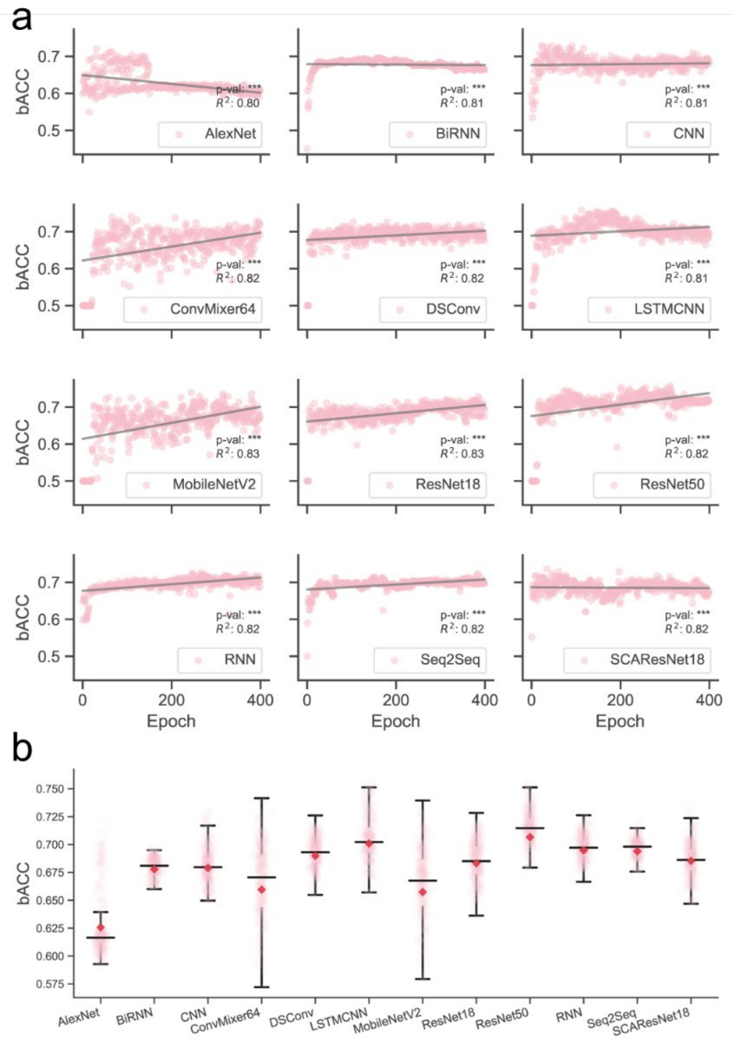


Figure S3. Full-scale bACC performance examination of deep learning algorithms on Test over training epochs. a. Landscapes of bACC variations. b. Boxplot of bACC values.

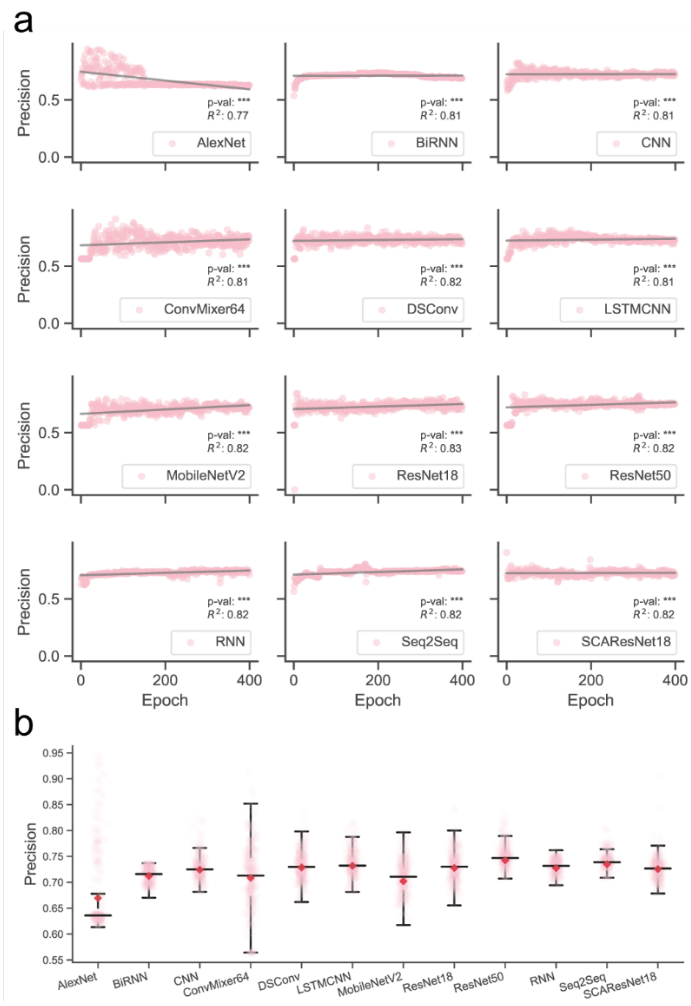


Figure S4. Full-scale precision performance examination of deep learning algorithms on Test over training epochs. a. Landscapes of precision variations. b. Boxplot of precision values.

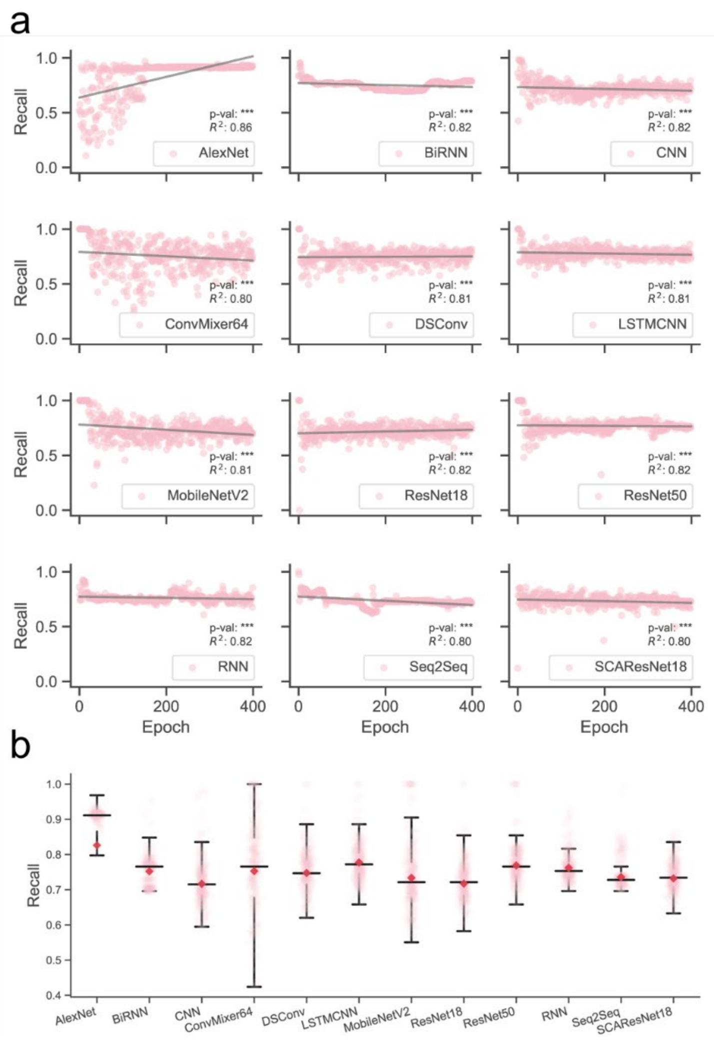


Figure S5. Full-scale recall performance examination of deep learning algorithms on Test over training epochs. a. Landscapes of recall variations. b. Boxplot of recall values.

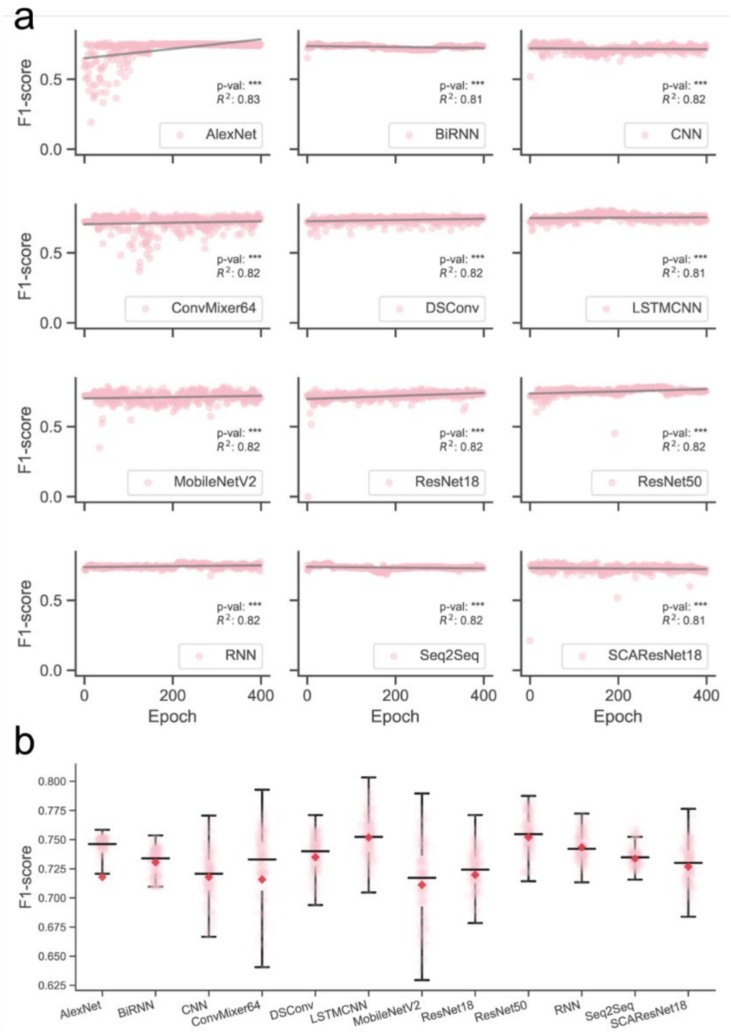


Figure S6. Full-scale F1-score performance examination of deep learning algorithms on Test over training epochs. a. Landscapes of F1-score variations. b. Boxplot of F1-score values.

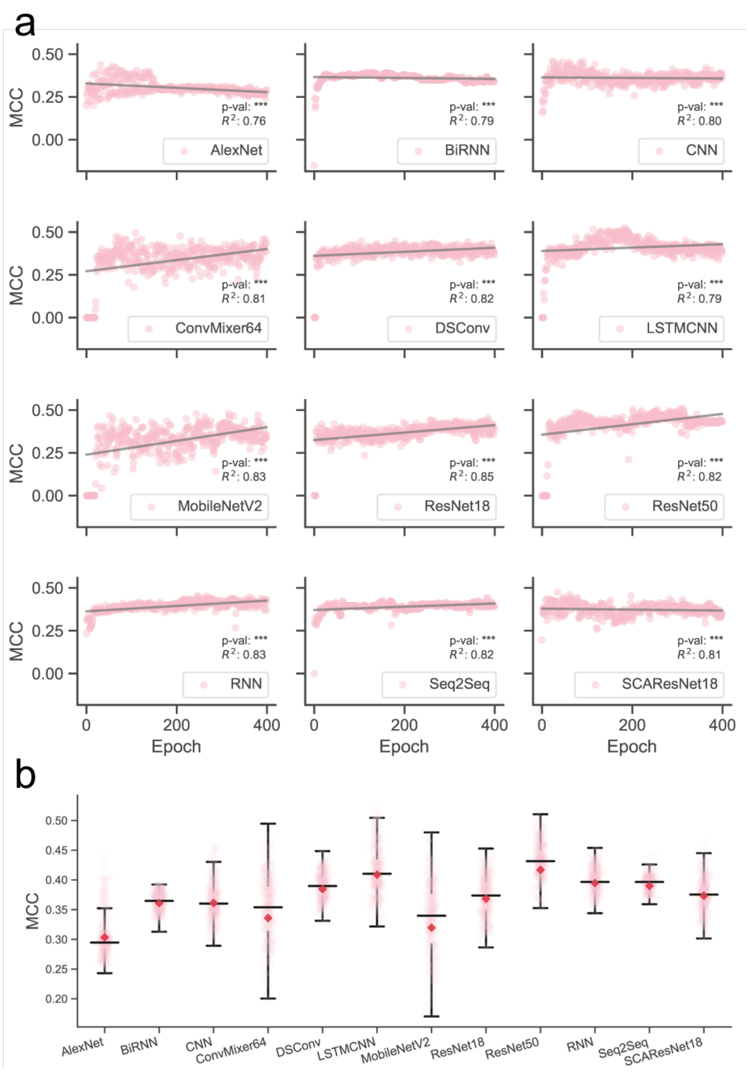


Figure S7. Full-scale MCC performance examination of deep learning algorithms on Test over training epochs. a. Landscapes of MCC variations. b. Boxplot of MCC values.

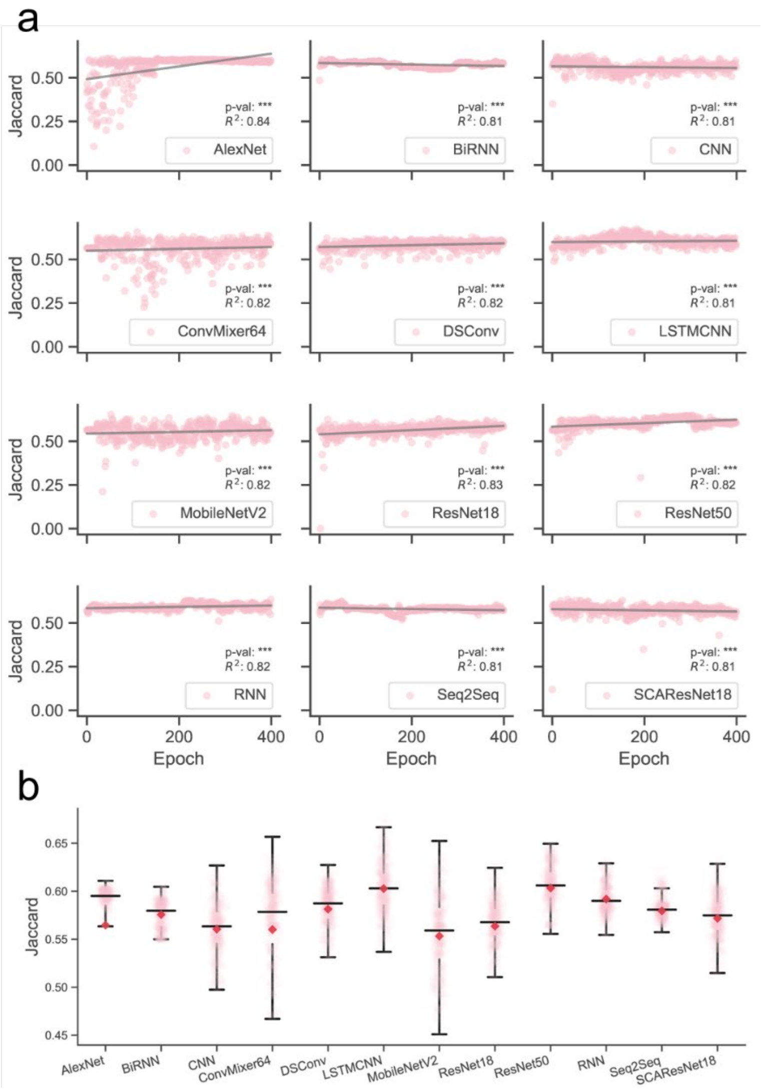


Figure S8. Full-scale Jaccard performance examination of deep learning algorithms on Test over training epochs. a. Landscapes of Jaccard variations. b. Boxplot of Jaccard values.

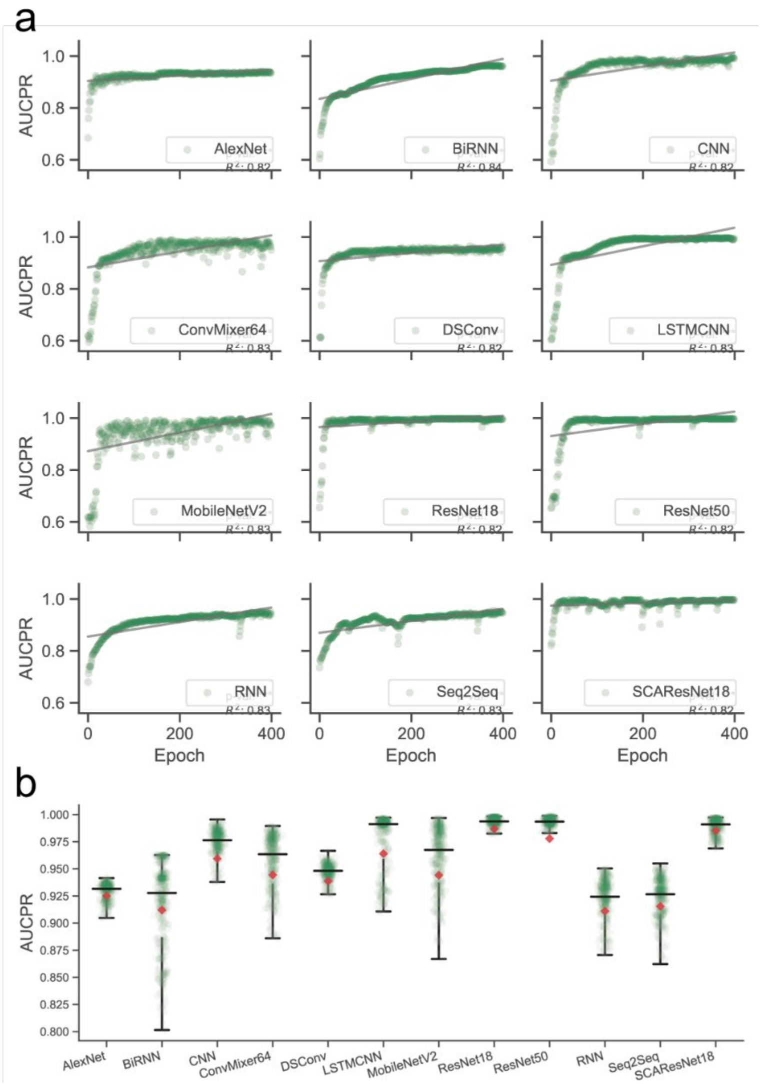


Figure S9. Full-scale AUCPR performance examination of deep learning algorithms on TestSim over training epochs. a. Landscapes of AUCPR variations. b. Boxplot of AUCPR values.

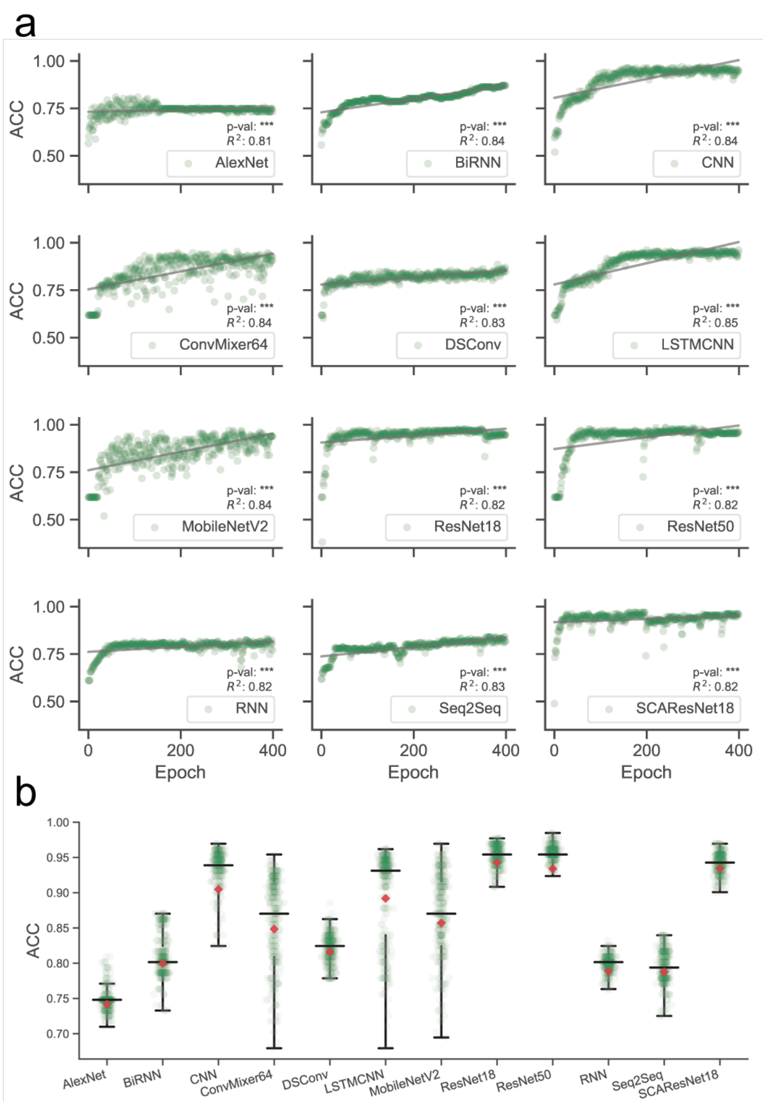


Figure S10. Full-scale ACC performance examination of deep learning algorithms on TestSim over training epochs. a. Landscapes of ACC variations. b. Boxplot of ACC values.

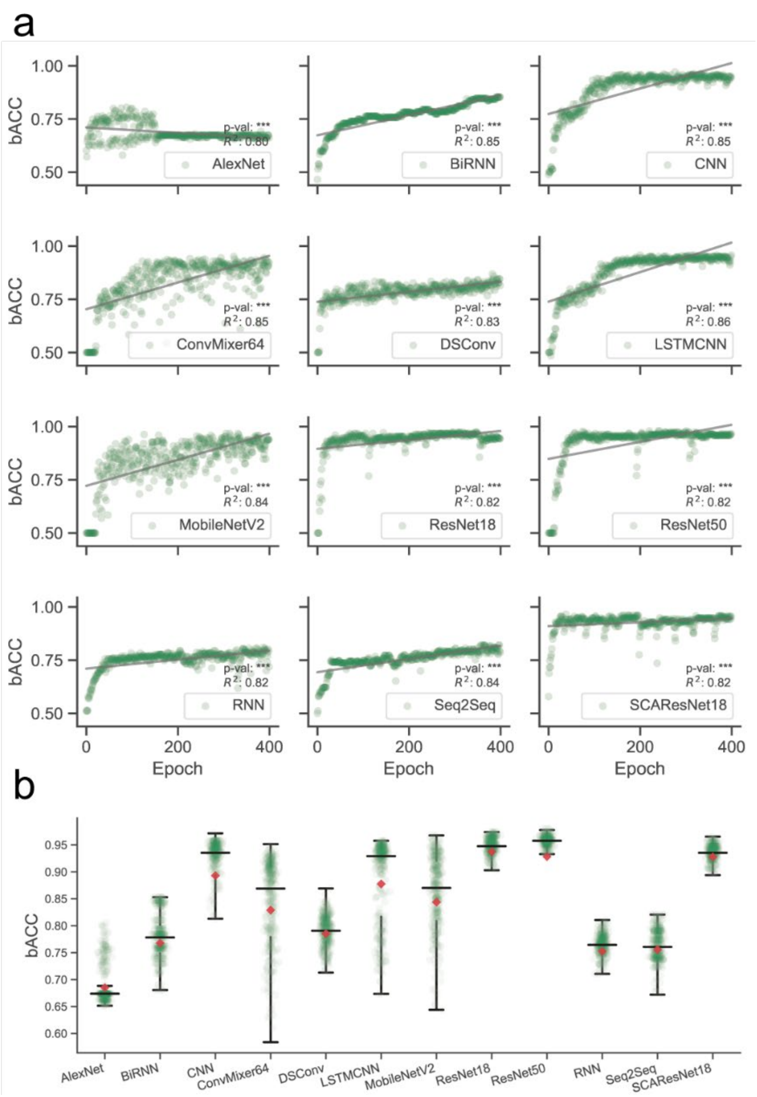


Figure S11. Full-scale bACC performance examination of deep learning algorithms on TestSim over training epochs. a. Landscapes of bACC variations. b. Boxplot of bACC values.

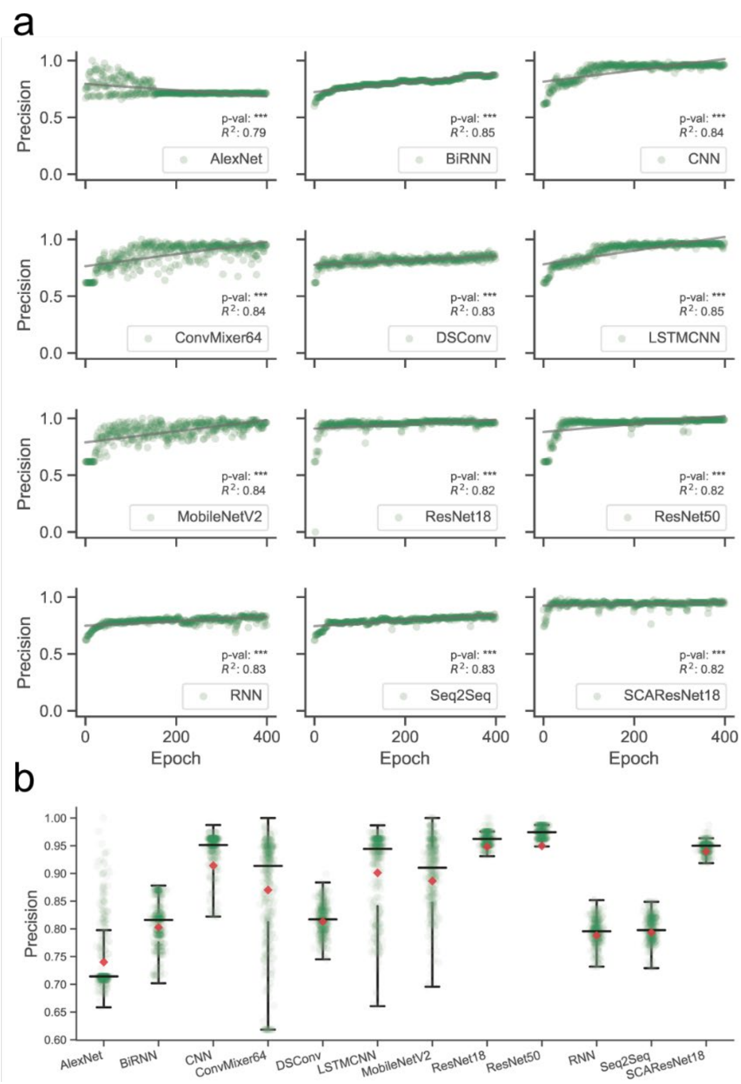


Figure S12. Full-scale precision performance examination of deep learning algorithms on TestSim over training epochs. a. Landscapes of precision variations. b. Boxplot of precision values.

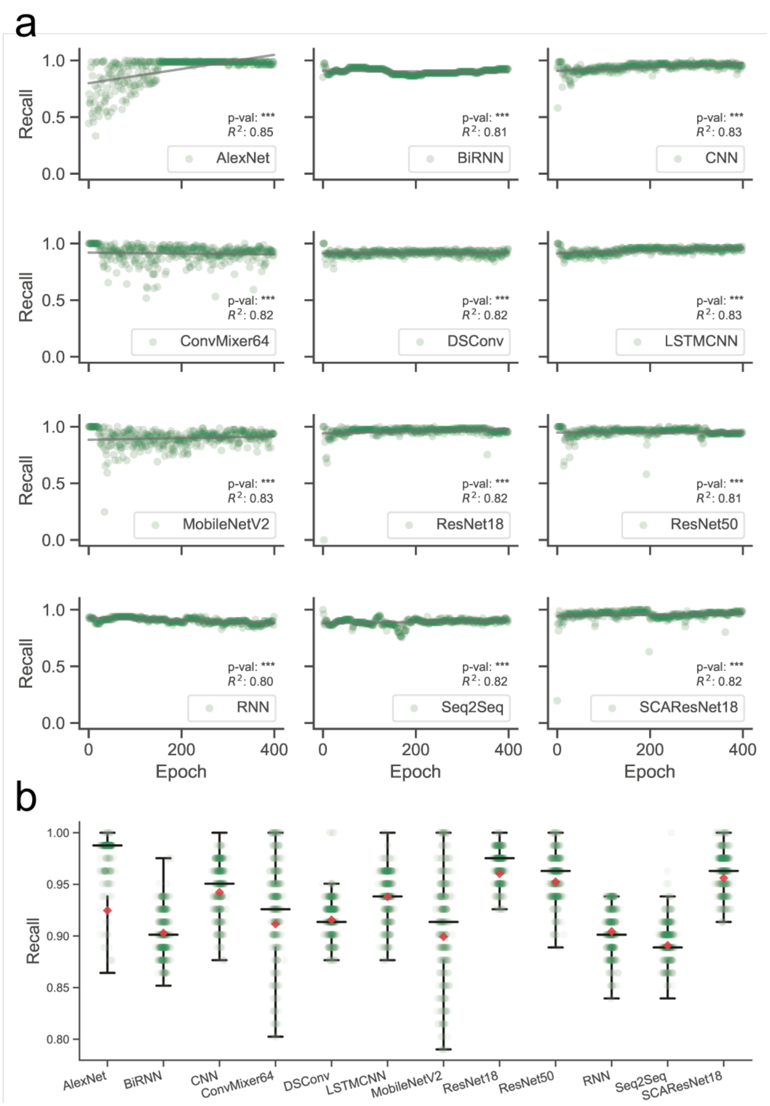


Figure S13. Full-scale recall performance examination of deep learning algorithms on TestSim over training epochs. a. Landscapes of recall variations. b. Boxplot of recall values.

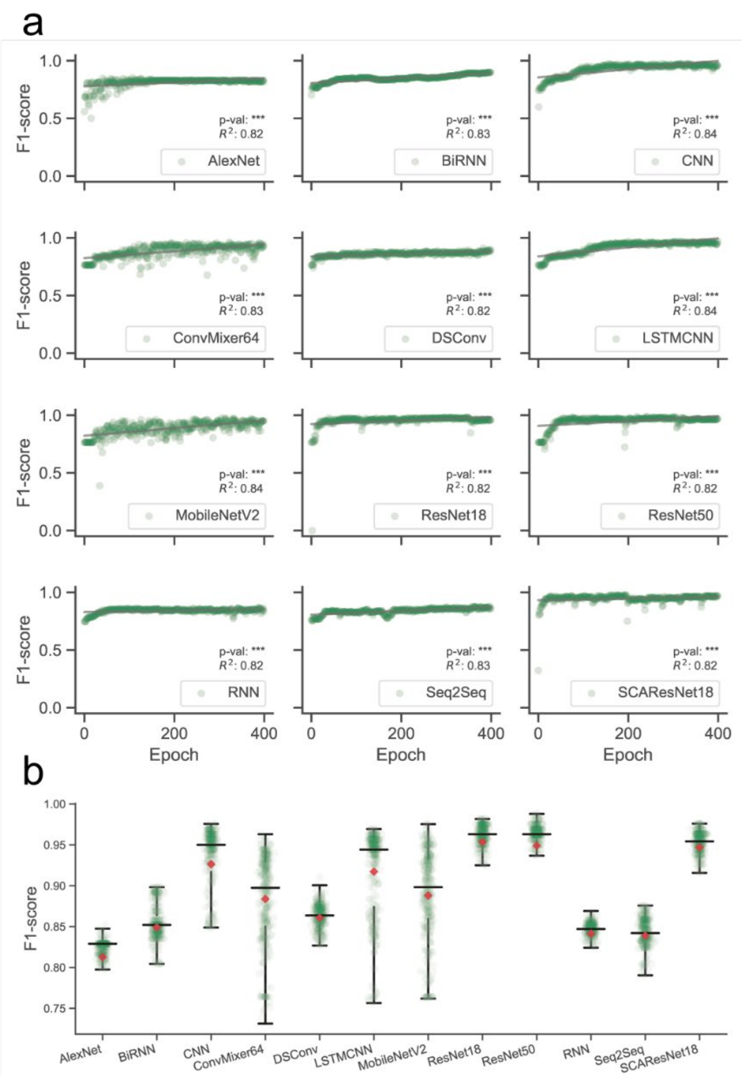


Figure S14. Full-scale F1-score performance examination of deep learning algorithms on TestSim over training epochs. a. Landscapes of F1-score variations. b. Boxplot of F1-score values.

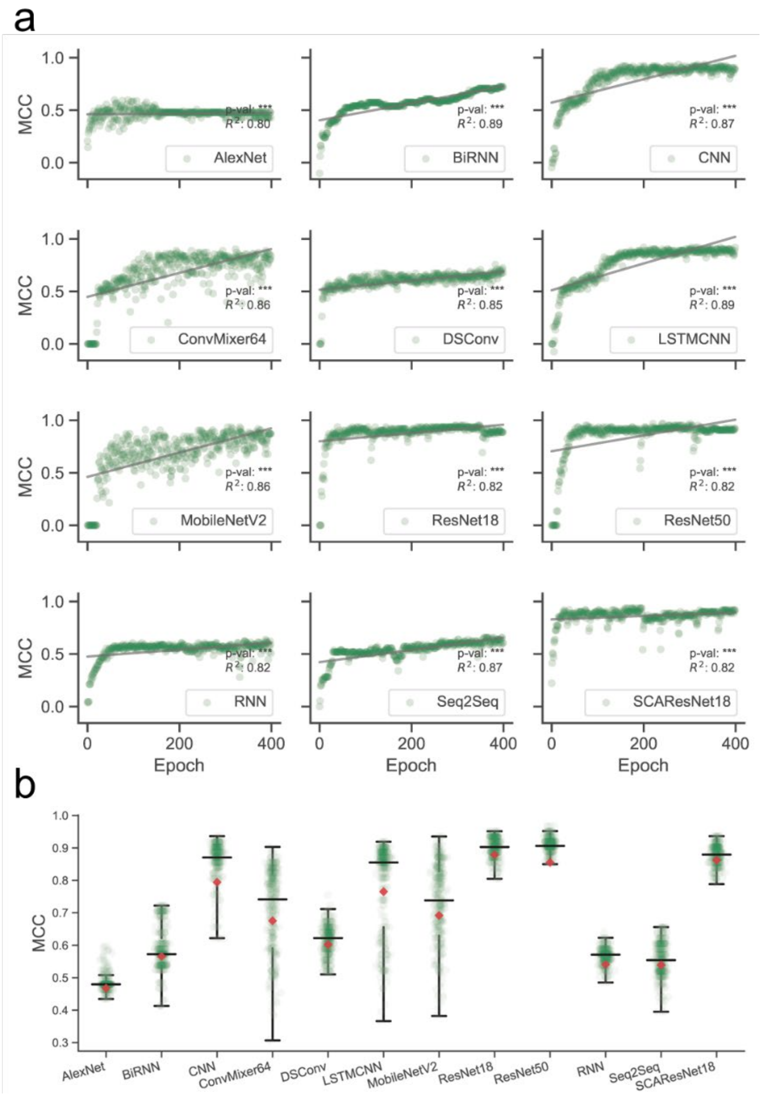


Figure S15. Full-scale MCC performance examination of deep learning algorithms on TestSim over training epochs. a. Landscapes of MCC variations. b. Boxplot of MCC values.

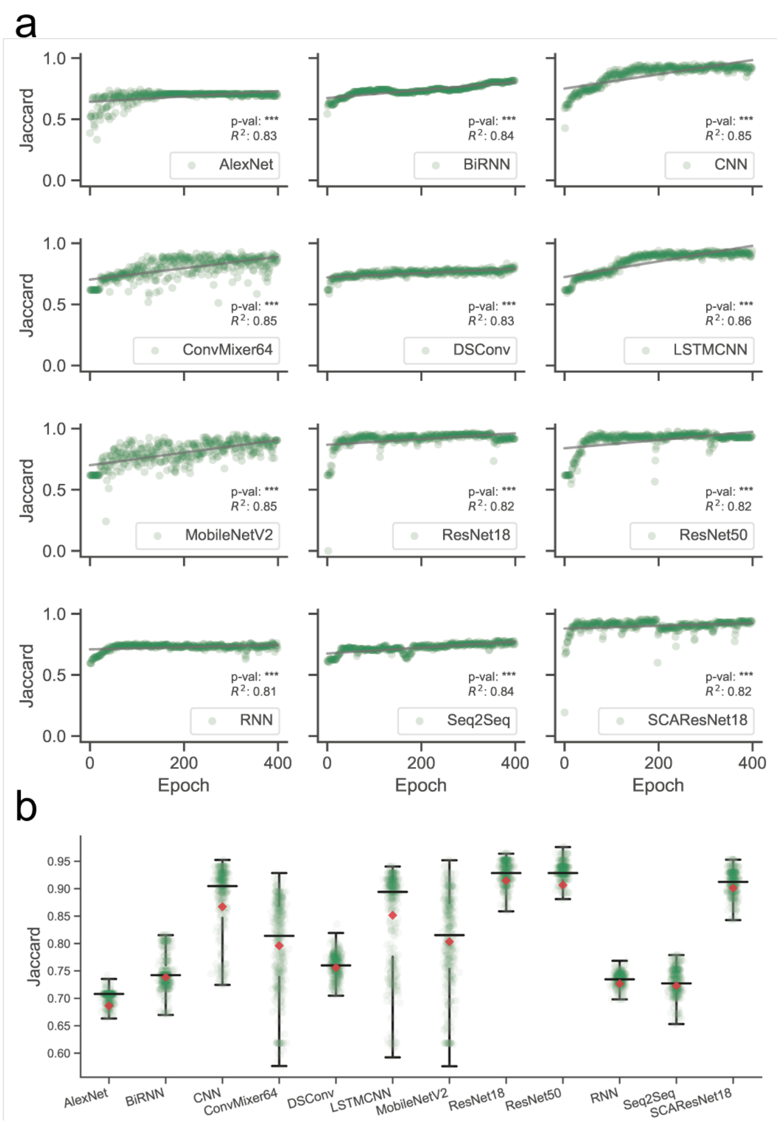


Figure S16. Full-scale Jaccard performance examination of deep learning algorithms on TestSim over training epochs. a. Landscapes of Jaccard variations. b. Boxplot of Jaccard values.

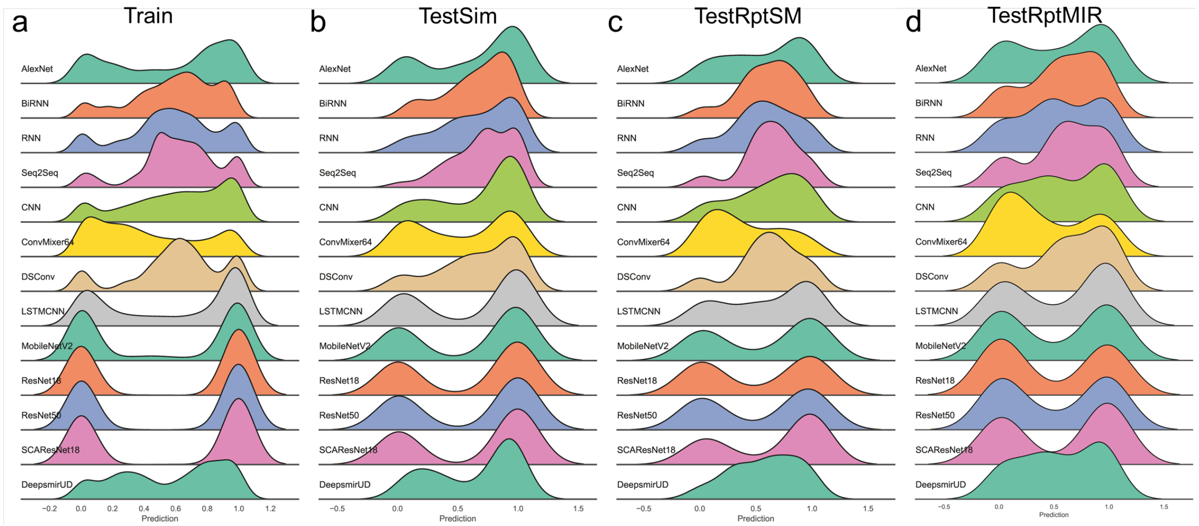


Figure S17. Ridge plot of prediction values (i.e., regulatory effects) on Train, TestSim, TestRptSM, and TestRptMIR.

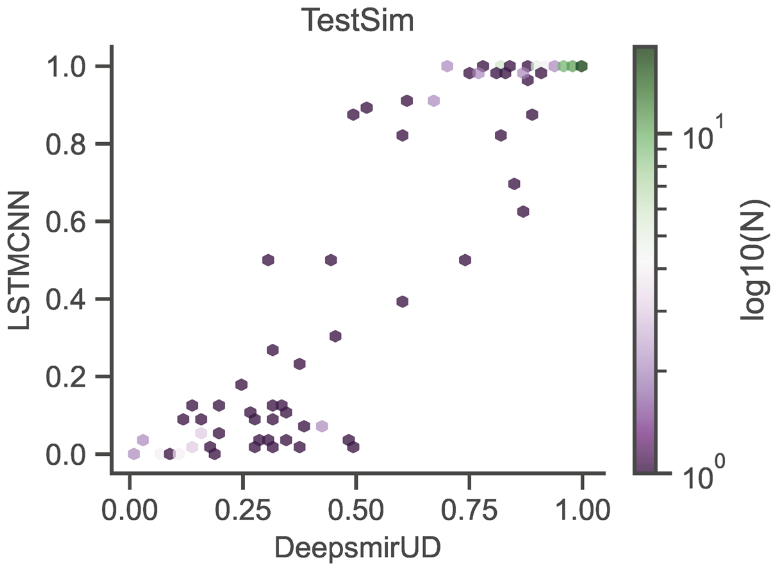


Figure S18. Hexagonal binned plot of comparison between LSTMCNN and DeepsmirUD predictions on TestSim.
